# A NRF2/β3-adrenoreceptor axis drives a sustained antioxidant and metabolic rewiring through the pentose-phosphate pathway to alleviate cardiac stress

**DOI:** 10.1101/2023.11.10.564735

**Authors:** Lauriane Y. M. Michel, Hrag Esfahani, Roxane Verdoy, Delphine de Mulder, Jérôme Ambroise, Véronique Roelants, Bertrand Bouchard, Jérôme Savary, Joseph P. Dewulf, Thomas Doumont, Caroline Bouzin, Vincent Haufroid, Joost J.F.P. Luiken, Miranda Nabben, Michael L. Singleton, Luc Bertrand, Matthieu Ruiz, Christine Des Rosiers, Jean-Luc Balligand

## Abstract

**Background:** Cardiac β3-adrenergic receptors (β3AR) are upregulated in diseased hearts and mediate antithetic effects to those of β1AR and β2AR. β3AR agonists were recently shown to protect from myocardial remodeling in preclinical studies and to improve systolic function in patients with severe heart failure. The underlying mechanisms, however, remain elusive.

**Methods:** To dissect functional, transcriptional and metabolic effects, hearts and isolated ventricular myocytes from mice harboring a moderate, cardiac-specific expression of a human *ADRB3* transgene (β3AR-Tg) and subjected to transverse aortic constriction (TAC) were assessed using echocardiography, RNAseq, PET scan, metabolomics, seahorse and metabolic flux analysis. Subsequently, signaling and metabolic pathways were investigated further *in vivo* in β3AR-Tg and *in vitro* in neonatal rat ventricular myocytes adenovirally infected to express β3AR and subjected to neurohormonal stress. These results were completed with an analysis of single nucleus RNAseq data from human cardiac myocytes from heart failure patients.

**Results:** Compared with WT littermate, β3AR-Tg mice were protected from hypertrophy after transaortic constriction (TAC), while systolic function was preserved. β3AR-expressing hearts displayed enhanced myocardial glucose uptake under stress in absence of increased lactate levels. Instead, metabolomic and metabolic flux analyses in stressed hearts revealed an increase in intermediates of the Pentose-Phosphate Pathway (PPP) in β3AR-Tg, an alternative route of glucose utilization, paralleled with increased transcript levels of NADPH-producing and rate-limiting enzymes of the PPP, without fueling the hexosamine metabolism. The ensuing increased content of NADPH and of reduced glutathione decreased myocyte oxidant stress, while downstream oxidative metabolism assessed by oxygen consumption was preserved with higher glucose oxidation in β3AR-Tg post-TAC compared to WT, together with increased mitochondrial biogenesis. Unbiased transcriptomics and pathway analysis identified NRF2 (NFE2L2) as upstream transcription factor which was functionally verified in β3AR-expressing cardiac myocytes where its translocation and nuclear activity was dependent on β3AR activation of nitric-oxide synthase (NOS) NO production.

**Conclusion:** Moderate expression of cardiac β3AR, at levels observed in human cardiac myocardium, exerts antioxidant effects through activation of the PPP and NRF2 pathway, thereby preserving myocardial oxidative metabolism, function and integrity under pathophysiological stress.

## Introduction

Myocardial hypertrophy is an essential part of adverse cardiac remodeling that occurs in response to pressure overload or neurohormonal stress and is a prequel to heart failure (HF). Hypertrophic remodeling contributes to lower ventricular compliance, to altered cardiac myocytes contractile properties and ultimately to the development of diastolic and/or systolic failure ^1, 2^. So far, very few therapeutic strategies proved to efficiently reverse cardiac hypertrophy or prevent the transition to heart failure.

In the course of hypertrophic remodeling, cardiac myocytes undergo an extensive gene expression reprogramming paralleled with increased levels of oxidative stress and metabolic remodeling ^3^. The latter is characterized by a loss of metabolic flexibility and a shift in substrate catabolism away from mitochondrial oxidation, increasing the reliance of the heart on glycolysis. ^4–6^. During this remodeling cardiac myocytes progressively become insensitive to insulin which further complicates the capacity for the heart to meet its energetic needs ^4–6^. In parallel, dysfunctional mitochondria contribute to the generation of intracellular reactive oxygen species (ROS) through electron leakage. Together with diminished endogenous antioxidant systems, these alterations elevate intracellular oxidative stress levels which further accentuate the cardiac hypertrophic response ^7, 8^.

We and others identified a protective action of cardiac β3AR against adverse cardiac remodeling ^9–11^. This beta3AR isotype has traditionally been considered as a “metabolic” receptor due to its well-characterized lipolytic and “beiging” effect in the adipose tissue ^12–14^. In the heart, β3AR expression, detected in human atrial and ventricular myocytes ^15^ is further upregulated in the diseased myocardium ^16, 17^ where β3AR activation alleviates oxidant stress and attenuates hypertrophy. While our previous work showed these protective effects to depend on β3AR activation of nitric oxide synthase (eNOS and nNOS) and cardiac NO production ^9–11^, the mechanistic link to the restoration of redox homeostasis was unknown. A recent cardiac proteomic screening in a systemic β3AR knock-out mouse model suggested an implication of β3AR in cardiac energy metabolism ^18^, but the underlying mechanism(s) remains elusive. Dissecting such mechanism(s) is particularly relevant given the therapeutic potential of new agonists of the human β3AR as suggested from recent clinical trials in heart failure ^19–21^.

The present work reveals the following major findings; we show that in the face of hemodynamic stress, expression of the human β3AR in cardiac myocytes increased insulin sensitivity and the uptake of glucose (but not fatty acids) in cardiac myocytes; that glucose is derived to the ancillary pentose phosphate pathway (PPP) with ensuing decreased intracellular oxidative stress levels and enhanced mitochondrial respiration; and that these effects are mediated by β3AR activation of the nuclear translocation of the transcription factor NRF2 (NFE2L2) through nitric oxide synthase (NOS).

## Methods

All data and materials are available within the article. Detailed methods are described in the Supplemental Material.

### Animal studies

All protocols were carried out in accordance with the Guide for the Care and Use of Laboratory Animals published by the U.S. National Institutes of Health (NIH) and the European Directive 2010/63/EU and were approved by local ethical committees.

### Data availability

Raw and processed RNA-seq data were deposited and made publicly available on the Gene Expression Omnibus (GSE230859). Single-nucleus RNA sequencing data were analyzed by using a recently published dataset generated from patients with heart failure who recovered (versus patients who did not recover) left ventricular systolic function after left ventricular assist device implantation^22^ (GSE226314).

### Statistical analysis

Results are expressed as mean ± SEM calculated from the average measurements obtained from each group of cells and mice. When normal distribution (tested by Kolmogorov-Smirnov test) was confirmed, raw data were analyzed using unpaired t-test or 2-way ANOVA followed by Sidak post-hoc test for multi-group comparisons. In absence of normal distribution, data were compared using nonparametric tests (Kruskall-Wallis followed by Dunn correction for multiple comparisons or Mann– Whitney). Statistical significance was accepted at the level of P<0,05.

## Results

### Anti-hypertrophic action of human β3-AR and protection against adverse remodeling

Adult mice heterozygous for cardiac-specific expression of human β3 adrenergic receptors (β3AR-Tg) ^10^ were subjected to transverse aortic constriction (TAC) and their cardiac phenotype characterized 9 weeks post-TAC and compared to control non-operated littermates. Note that in this model, the abundance of human β3AR proteins in heterozygous animals is comparable to that observed in human ventricular biopsies ^11^. Nine weeks post-surgery β3AR-Tg animals developed a significantly milder hypertrophic response than WT littermates as assessed by heart weight measurements (**Fig 1A-1B**). Trans-stenoticgradientswereassessedinallmicepost-surgerytoensurecomparablecardiac afterload between genotypes (**Fig 1C**) with a maximum velocity ranging from 2,5 m/s to 4,5 m/s. In this mouse strain, the relatively mild trans-stenotic gradients recapitulated the early stages of cardiac remodeling without established alterations of systolic function leading to heart failure (**Suppl. Table 1**). Accordingly, ejection fraction and fractional shortening were only slightly decreased in WT littermates and fully preserved in β3AR-Tg (**Suppl. Table 1; Suppl Fig. S1A-S1B;**). A clear attenuation of left ventricular mass was observed in β3AR-Tg mice (**Fig 1C-1D**) which was not at the expense of cardiac function as ejection fraction and fractional shortening were maintained to their levels before surgery as was the survival rate (**Suppl Fig S1A-S1B**). In addition to cardiac hypertrophy, a significant increase in circulating BNP was observed in WT post-TAC (**Fig 1F**), with levels substantially lower than observed in the failing hearts consistent with an early stage of remodeling. Nonetheless, markedly lower circulating BNP levels were observed in β3AR-Tg (**Fig. 1F**). In addition, lower hypertrophic response in β3AR-Tg post-TAC was confirmed histologically by myocyte transverse area (**Fig. 1G**). Similarly, adenoviral expression of the human β3AR (Ad-β3AR) in primary neonatal rat ventricular myocytes abrogated the hypertrophic response classically triggered under neurohormonal stress by phenylephrine and observed in control conditions (Ad-GFP) (**Fig 1H**). Finally, expression levels of hypertrophic markers (ANP, BNP, βMHC and βMHC/ αMHC) were sizeably lower in adult ventricular myocytes isolated from β3AR-Tg hearts post TAC compared to their WT littermates (**Fig 1I-1L**).

**Figure 1.**
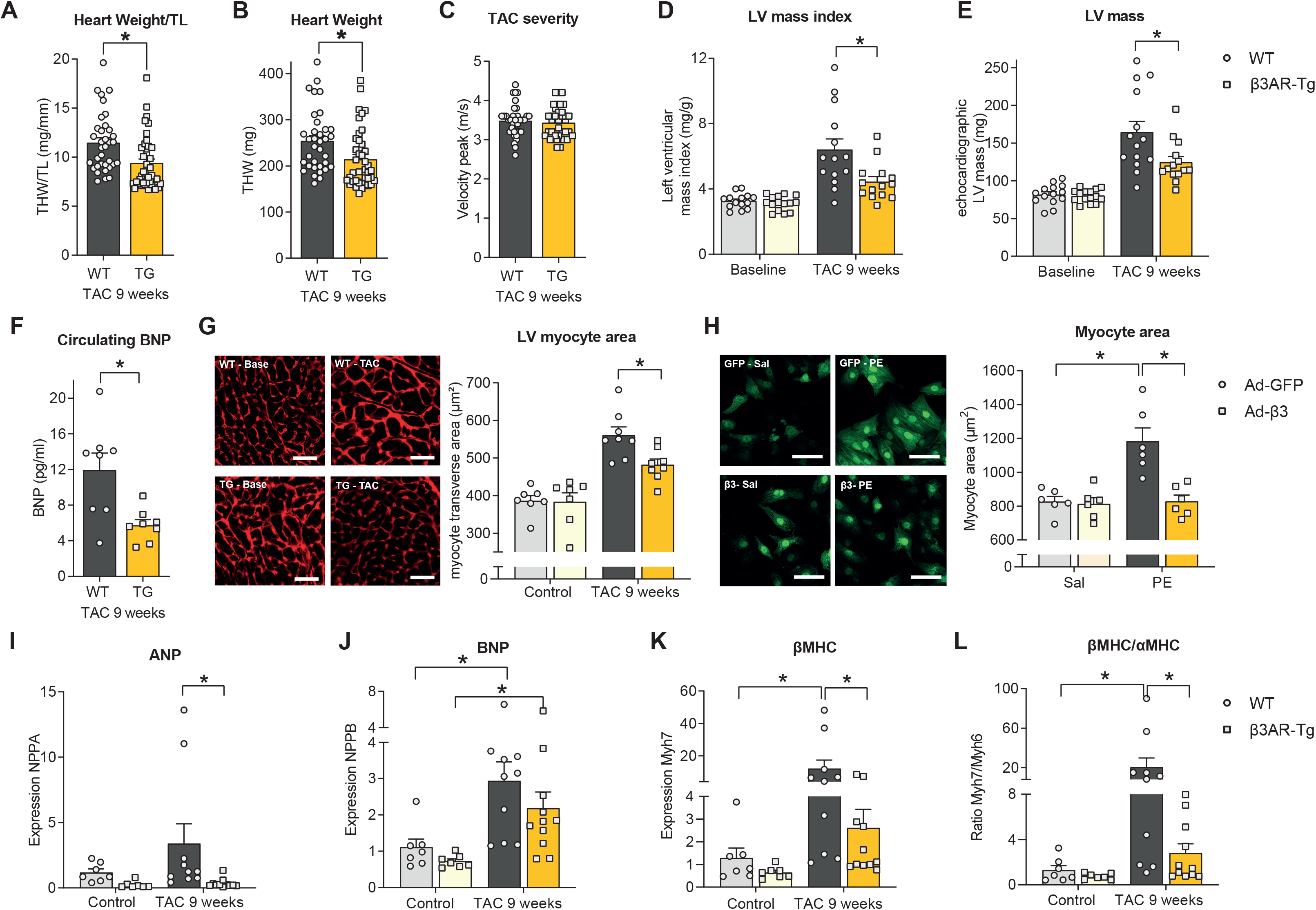
Cardiac expression of human β3AR attenuates myocardial hypertrophy and increases glucose entry under hemodynamic stress. – (A) Total heart weight normalized on tibial length (TL) and (B) Total heart weight for all animals subjected to transverse aortic constriction (TAC) and associated (B) trans-stenotic gradient severity (by echocardiography) following surgery associated. (D-E) Echocardiographic follow-up of left ventricular mass index and left ventricular mass before and 9 weeks after TAC surgery. (F) Plasma BNP levels post-TAC. (G) Representative images and quantification of myocyte area measured histologically (WGA staining; each dot is mean from >200 myocytes per heart) in left ventricular transverse sections from β3AR-Tg and WT littermates subjected or not to TAC (scale 20µm). (D) Left ventricular ejection fraction before and 9 weeks after TAC surgery. (H) Representative images and quantification of myocyte area in neonatal rat ventricular myocytes (NRVM) infected with Ad-GFP-β3AR (β3AR) or Ad-GFP (GFP) treated with Phenylephrine (PE, 50µM) or Saline (Sal)(scale 20µm). (I-L) Hypertrophic transcriptional program assessed by (I) ANP (J) BNP (K) βMHC transcripts abundance and (L) associated βMHC/αMHC ratio in AVM isolated from 9-week TAC or Control β3AR-Tg or wild-type mice. Results are expressed as means ± SEM with *P < 0,05 calculated by Two-way Anova and corrected for multiple comparisons with Sidak’s test, except for A, B, C and F analyzed by unpaired t-test and (D-E) analyzed by mixed models corrected with Sidak’s test.

### Increased glucose uptake and attenuated hypertrophy by cardiac β3AR upon hemodynamic stress

We next tested whether the protection mediated by β3AR was associated with changes in myocardial glucose or lipid handling. With this aim, β3AR-Tg mice were subjected to intraperitoneal glucose and insulin tolerance test (GTT and ITT, **Fig 2A-B** respectively) before and after aortic constriction. The normalization of glycemic profile was unchanged among genotypes after challenging the animals with glucose 2g/kg (**Fig 2A**). Strikingly, insulin administration (0,5U/kg) led to a lower glycemia in β3AR-Tg 9 weeks post-TAC compared to controls suggesting an improved glucose uptake by peripheral tissues in β3AR-Tg (**Fig 2B**); an effect not detectable during GTT due to lower circulating insulin necessary to normalize glycemia in β3AR-Tg as measured during glucose administration (**Fig 2C**). As the human β3AR transgene expression under the *Myh6*promoteris restricted to cardiac myocytes, this effect could be attributed either to an enhanced glucose uptake capacity specific to the cardiac myocytes or to factor(s) released by β3AR-Tg myocytes affecting peripheral tissues. Unbiased serum metabolomics did identify a number of enriched metabolites after TAC compared to baseline (**Suppl Fig S2A**), but these did not differ between β3AR-Tg and WT post-TAC (**Fig 2D** **and Suppl FigS2B**) or in baseline condition (**Suppl Fig S2C**) suggesting a myocyte-specific effect rather than a peripheral effect. Therefore, we next compared the glucose uptake capacity of the heart *in vivo* between the two genotypes. ^18^F-fluorodeoxyglucose (^18^FDG) uptake confirmed an increase in myocardial glucose uptake in β3AR-Tg 9 weeks post-TAC (**Fig 2E**). In isolated adult ventricular myocytes (AVM) from similarly treated animals, insulin-dependent stimulation of tritiated glucose uptake was reduced in WT, but restored in β3AR-Tg (**Fig 2F**). Accordingly, expression of the insulin-dependent glucose transporter Glut4 was preserved after TAC in β3AR AVM, but downregulated in WT AVM, while expression levels of glucose transporter Glut1 were unchanged (**Fig 2G-H**). Conversely, myocardial fatty acid uptake assessed *in vivo* by 18-^18^F-Fluoro-4-Thia-Oleate (**Suppl Fig S3A**) and by ^14^C-Palmitate in AVM was unchanged (**Suppl Fig S3B-C)** as was the expression level of fatty acid transporter CD36 (**SupplFigS3D**). Therefore, moderate expression of β3AR in cardiac myocytes increases insulin sensitivity, glucose (but not lipid) uptake and attenuates hypertrophy upon chronic hemodynamic overload.

**Figure 2.**
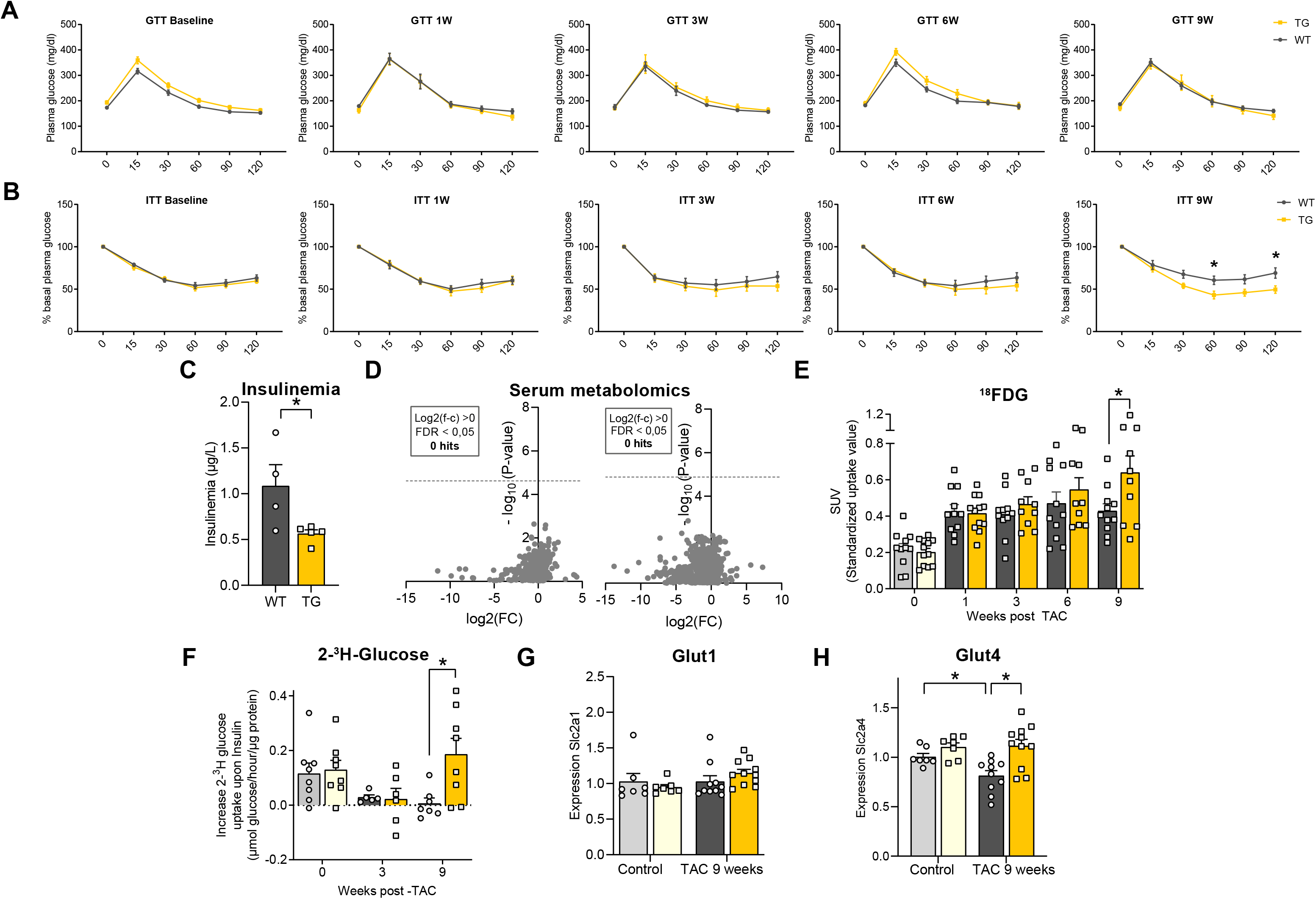
Cardiac expression of human β3AR increases glucose entry and insulin sensitivity under hemodynamic stress. **– (A-B)** Intraperitoneal **(A)** Glucose (2g/kg) and **(B)** Insulin tolerance test (0,5U/kg) before and over time up to 9 weeks following aortic constriction in β3AR-Tg and WT mice. **(C)** Insulinemia measured in serum from β3AR-Tg and WT animals 9 weeks post-TAC measured 30 minutes after intraperitoneal glucose injection. **(D)** Volcano plot displaying differential unbiased serum metabolomics from 9-weeks TAC β3AR-Tg compared to 9-weeks TAC WT mice analyzed by reverse phase LC/MS in negative (left) and positive (right) ionization mode (1886 and 3881 detected and plotted metabolites, respectively; see Methods for details). **(E)**Myocardial ^18^F-fluorodeoxyglucose (^18^FDG) uptake measured by Positron Emission Tomography (PET) *in vivo* (expressed as standardized uptake value) and **(F)** Increase in tritiated glucose uptake following insulin-treatment in adult ventricular myocytes (AVM) from β3AR-Tg or wild-type littermates before surgery or at different time points post-TAC. **(J-K)** glucose transporters **(J)** Glut 1 and **(K)** Glut 4 transcripts abundance in AVM isolated from 9-week TAC or Control β3AR-Tg or wild-type mice. Results are expressed as means ± SEM with *P < 0,05 calculated by Two-way Anova and corrected for multiple comparisons with Sidak’s test, except for C analyzed by unpaired t-test.

### Increased glucose uptake and glycogenolysis by cardiac β3AR fuel the pentose phosphate pathway

The intracellular fate of glucose was further explored. In addition to glucose entry as glucose-6-phosphate (**Fig 2E**), initial targeted metabolomic analyses revealed a marked decrease in the upper glycolytic intermediates directly downstream of fructose-6-phosphate (F1,6P and DHAP; **Fig 3B**) in β3AR-Tg hearts post-TAC indicating an escape route from the classical glycolysis. UDP-GlcNAc levels were unaffected by hemodynamic stress or genotypes eliminating an enrichment of the hexosamine biosynthetic pathway (**Fig 3C**). UDP-Glucose, an intermediate of glycogenesis was unchanged in β3AR-Tg hearts, arguing against derivation towards glycogen synthesis (**Fig 3D**). Instead, glycogen content was markedly reduced in β3AR-Tg hearts (**Fig 3E**), paralleled with an increase in expression levels of Phosphorylase Kinase Regulatory Subunit Alpha (PHKA) and gamma (PHKG), key enzymes of glycogenolysis (**Fig 3F-3G**), likely further fueling the glucose supply upstream from glycolysis. Evaluation of the escape towards the pentose phosphate pathway (PPP) revealed an enrichment in all detectable intermediates i.e. ribose-5-phosphate and sedoheptulose-7-phosphate (**Fig 3H-I**), further supporting an escape route from the classical glycolysis towards the PPP. Fluxes through the upper glycolysis and the PPP were further dissected by metabolic flux analysis on Langendorff-perfused hearts from 9-week TAC mice perfused with uniformly labelled U-^13^C_6_-Glucose followed by LC-MS (**Fig 4A**). This revealed a significant enrichment of the upper glycolysis intermediates in β3AR-Tg hearts (vs. WT) at short perfusion times, followed by a plateau of maximal ^13^C molar percent enrichment in both genotypes (**Fig 4A-B**). Coincidentally, a progressive enrichment over time of the intermediates of the PPP was observed in both genotypes, among which Ribose-5P (R5P) and sedoheptulose-7P (S7P) were increased in β3AR-Tg hearts (vs. WT) (**Fig 4C-D**). Conversely, there was no enrichment in UDP-GlcNAc, arguing against glucose metabolic redirection towards the Hexosamine biosynthetic pathway (**Fig 4E**). Examination of the differential expression of key enzymes in the PPP between AVM from 9-week TAC β3AR and WT mice revealed increased transcripts abundance of the NADPH-producing enzymes, Phosphogluconate dehydrogenase (PGD) and Glucose-6-phosphate dehydrogenase (G6PD) in the oxidative phase of the pathway (**Fig 4F-G**) as well as Transaldolase (Taldo) in the lower PPP (**Fig 4H**).

**Figure 3.**
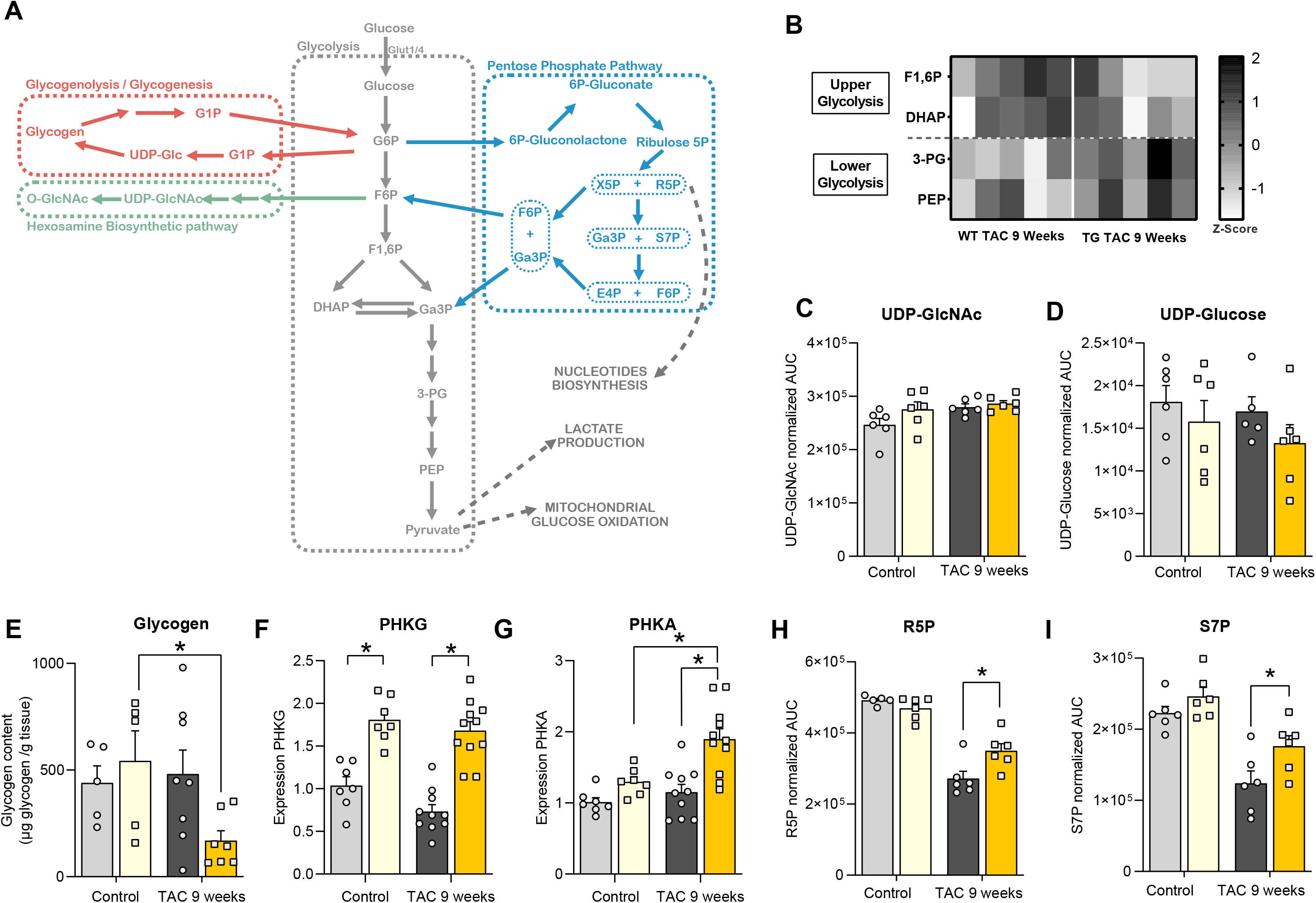
Myocardial glucose is rerouted away from classical glycolysis by β3AR. **– (A)** Schematic representation of the glucose metabolic pathways and their interactions **(B-I)** Targeted metabolomics measurements of intermediates of glycolysis and accessory pathways by LC/MS in cardiac extracts from β3AR-Tg and WT mice at baseline and 9 weeks after TAC surgery. **(B)** Heatmap of Z-score of glycolysis intermediates **(C)** UDP-GlcNAc and **(D)** UDP-Glucose **(H)** Ribose-5-phosphate **(I)** Sedoheptulose-7-phosphate. **(E)** Quantitation of glycogen content in cardiac extracts; **(F-G)** glycogenolysis enzymes: **(F)** Phosphorylase Kinase Regulatory Subunit Gamma (PHKG) **(G)** and alpha (PHKA) transcripts abundance in AVM isolated from 9-week TAC or Control β3AR-Tg or wild-type mice mice. Results are expressed as means ± SEM with *P < 0,05 calculated by Two-way Anova corrected for multiple comparisons with Sidak’s test

**Figure 4.**
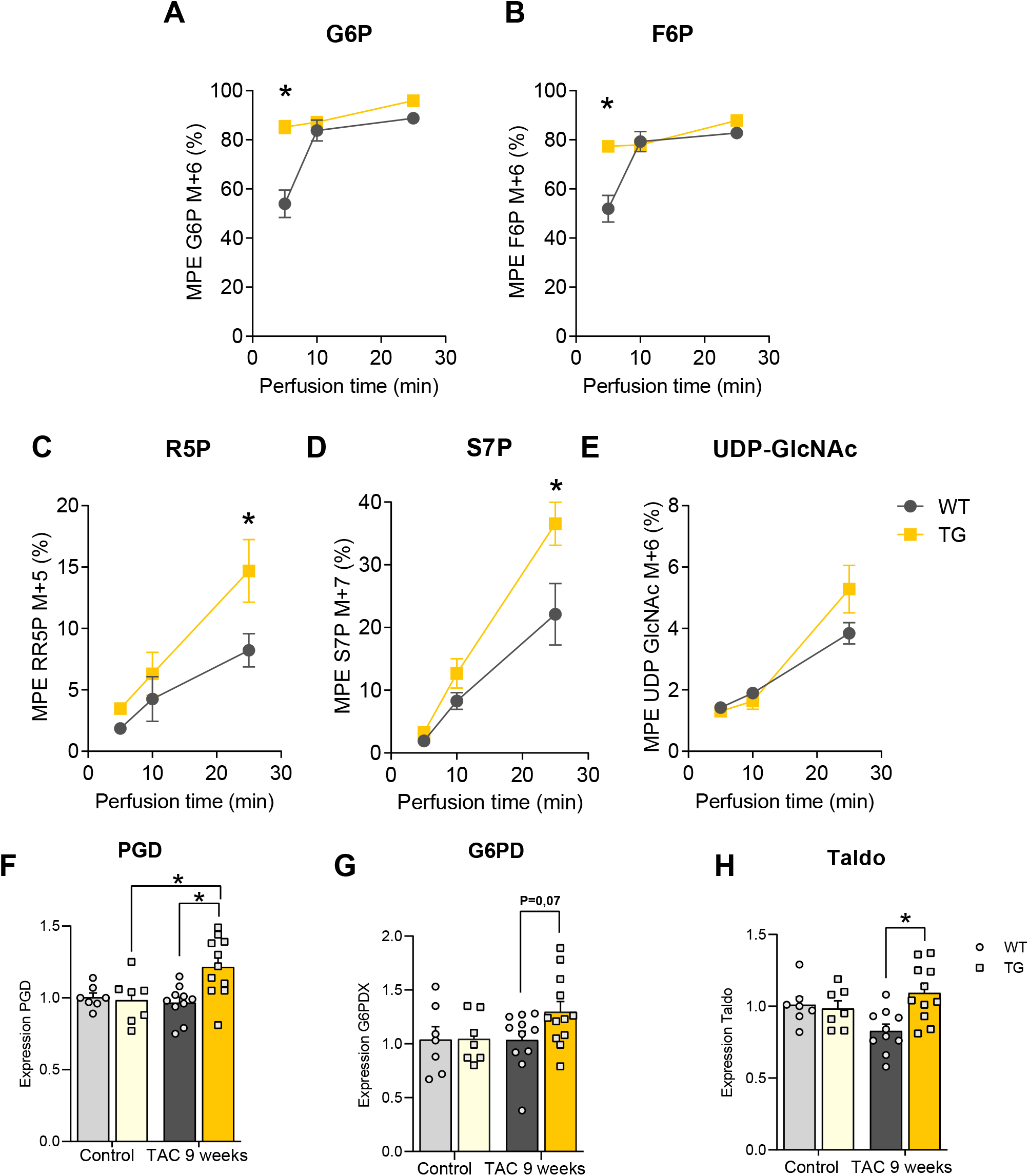
Myocardial glucose is rerouted to fuel the pentose phosphate pathway. **–(A-E)** Molar percent enrichment of [U-^13^C_6_] in upper glycolysis and pentose phosphate pathway intermediates as a function of perfusion duration with [U-^13^C_6_]-Glucose in Langendorff-perfused hearts from β3AR-Tg and WT mice 9 weeks post-TAC. **(A)** Glucose-6-phosphate M+6 **(B)** Fructose-6-phosphate M+6 **(C)** UDP-Glc-NAc M+6 **(D)** Ribose/Ribulose-5-Phosphate M+5 **(E)** Sedoheptulose -7-phosphate M+7. **(F-H)** Pentose phosphate pathway enzymes: **(F)** Phosphogluconate dehydrogenase (PGD) **(G)** Glucose-6-phosphate dehydrogenase (G6PD) and **(H)** Transaldolase (Taldo) transcripts abundance in AVM isolated from 9-week TAC β3AR-Tg or wild-type mice.

### Antioxidant effect of cardiac β3AR through an improved NADPH-mediated cellular detoxification

Having demonstrated the increased flux through the pentose phosphate pathway, we next assessed the consequences of the upregulation of PPP enzymes, specifically the NADPH-producing enzymes present in the oxidative phase of the pathway (**Fig 5A**). NADPH levels were considerably higher in 9-week TAC β3AR-Tg compared to WT (**Fig 5B**) whereas NADP^+^ levels were unmodified (**Fig 5C**) leading to a NADP ^+^/NADPH ratio maintained at a low level in β3AR-Tg compared to the unbalanced ratio observed in WT hearts (**Fig 5D**). NADPH being instrumental to reduce the pool of oxidized glutathione (GSSG) during cellular detoxification, we evaluated glutathione availability. Reduced glutathione was higher in 9-week TAC β3AR-Tg hearts suggesting an improved redox turnover of glutathione (**Fig 5E**), as confirmed by the maintained GSH/GSSG ratio (**Fig 5F**). Together with upregulated transcripts of glutathione peroxidase Gpx1 (**Fig 5G**), a detoxification enzyme responsible for scavenging reactive oxygen species (ROS), these data pointed towards increased antioxidant capacity of β3AR-expressing hearts under stress. Accordingly, the abundance of the oxidative stress marker Methionine sulfoxide (MetO) was reduced in β3AR-Tg hearts (**Fig 5H**). Finally, measurements of ROS levels in AVM freshly isolated from β3AR-Tg and WT hearts showed a significant decrease in intracellular ROS in β3AR-expressing cardiac myocytes in basal conditions (**Fig 5I**) or after exposure to 20µM H _2_O_2_ (**Fig 5J**). Therefore, the combined observations of increased antioxidant capacity in stressed hearts all support an increased flux of the oxidative PPP in β3AR-Tg.

**Figure 5.**
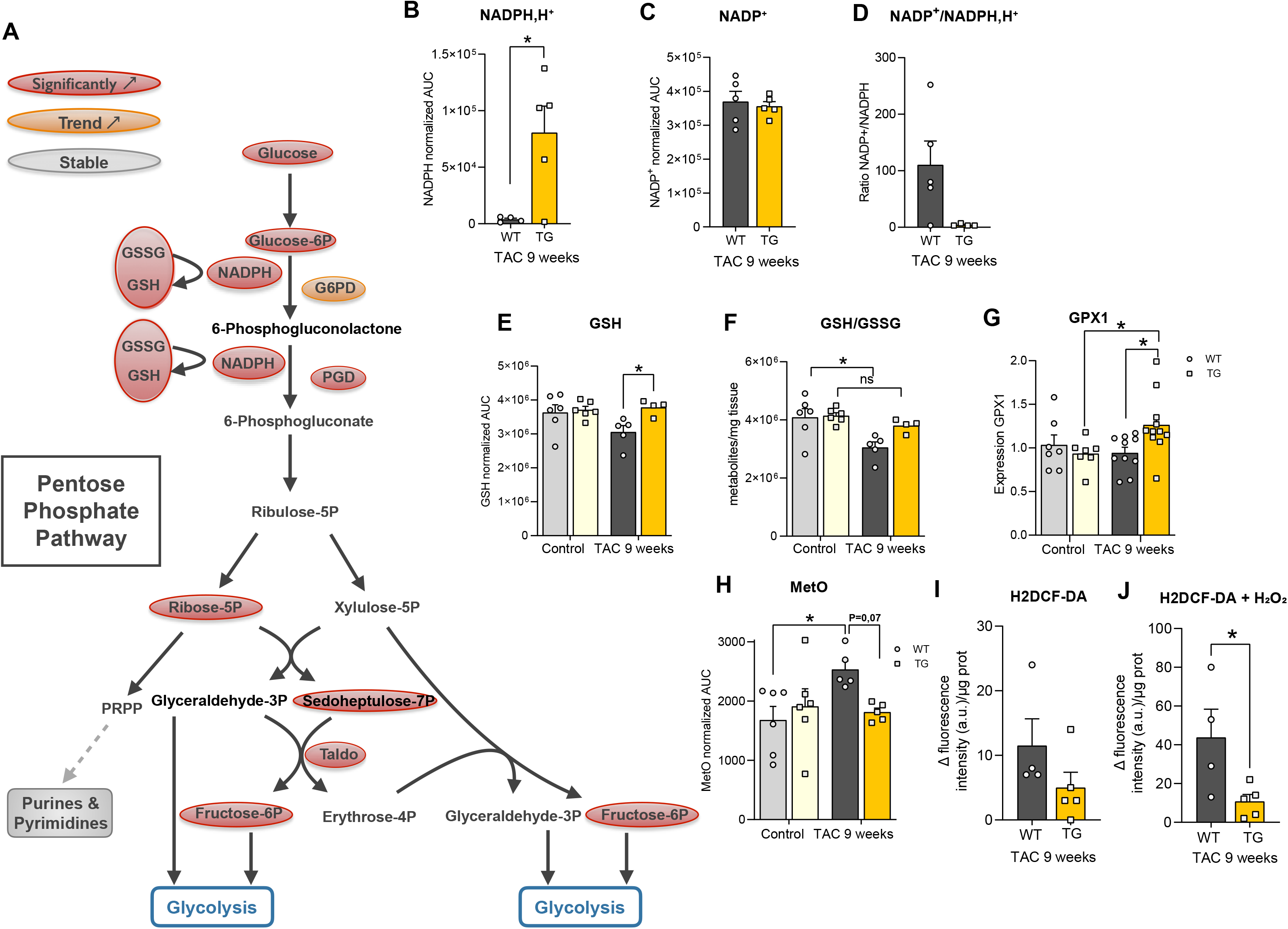
PPP protects against oxidative stress post-TAC through an improved NADPH-mediated cellular detoxification. **– (A)** Reaction steps of the pentose phosphate pathway highlighting the β3AR-mediated significant increases (red), trend for increase (orange) and no change (grey) in metabolic intermediates. **(B-D)** LC/MS determination of **(B)** NADPH levels **(C)** NADP ^+^ levels and **(D)** corresponding NADP+/NADPH ratio in cardiac extracts from β3AR-Tg and WT mice 9 weeks post-TAC determined by HILIC LC/MS. **(E)** quantification of reduced glutathione (GSH) and **(F)** reduced over oxidized glutathione ratio in cardiac extracts from 9-week TAC or Control β3AR-Tg or wild-type mice. **(G)** glutathione peroxidase 1 (Gpx1) transcript abundance in AVM isolated from 9-week TAC or Control β3AR-Tg or wild-type mice. **(H)** LC/MS quantification of methionine sulfoxide (MetO) in cardiac extracts from 9-week TAC or Control β3AR-Tg or wild-type mice **. (I-J)** Comparative increase in intracellular reactive oxidant species (ROS) measured by H _2_DCFDA fluorescence in isolated AVM from β3AR-Tg and WT hearts 9 weeks post-TAC **(I)** in basal conditions and **(J)** during exposure to extracellular H_2_O_2_ (20µM) (see methods for details). Results are expressed as means ± SEM and **(E-H)** analyzed by Two-way Anova corrected for multiple comparisons with Sidak’s test and **(B-D & H-J)** analyzed by unpaired t-test ; *P < 0,05.

### Cardiac β3AR feeds lower glycolysis through PPP intermediates and supports mitochondrial respiration

AsintermediatesofthePPPcanfeedthebiosynthesisofnucleotidesfromribose-5-phosphate (**Fig 5A**), we compared levels of purines from our targeted metabolomic data and found similar levels among genotypes (**Fig 6A**) rather suggesting a re-routing of non-oxidative PPP intermediates back into glycolysis. This was also consistent with increased abundance of lower glycolysis intermediates, such as 3-Phosphoglycerate (3-PG) and Phosphoenolpyruvate (PEP) (**Fig 3B**). Metabolic flux analysis of effluents from 9-week TAC perfused hearts labelled with U-^13^C_6_-Glucose and analyzed by GC-MS showed an enrichment of labelled pyruvate (**Fig 6B**) but not labelled lactate (**Fig 6C**) in β3AR-Tg hearts; similarly, cardiac release rate of lactate, reflecting its formation through glycolysis was not increased in β3AR-Tg hearts (**Fig 6D**) and extracellular acidification resulting from the proton release associated with lactate production was not increased in AVM from the same β3AR-Tg hearts (**Fig Suppl S4**). Instead β3AR expression promoted O _2_ consumption, as measured in Langendorff-perfused 9-week TAC hearts (**Fig 6E**) as well as in AVM from similar hearts, in which β3AR expression enabled the recovery of maximal respiration capacity as recorded by Seahorse analyzer (**Fig 6F-G**); thus pointing towards an accrued mitochondrial oxidation, particularly of glucose, as substrate complementation in the Seahorse media was restricted solely to glucose. In addition, transcript abundance of factors implicated in mitochondrial biogenesis (PGC1α, TFAM) and mitochondrial fusion (Mitofusin 1, Mitofusin 2, Opa 1) (**Fig 6H**) were upregulated in 9-week TAC β3AR-Tg hearts. Therefore, cardiac β3AR re-derives intermediates of the PPP towards lower glycolysis while preserving mitochondrial oxidative metabolism.

**Figure 6.**
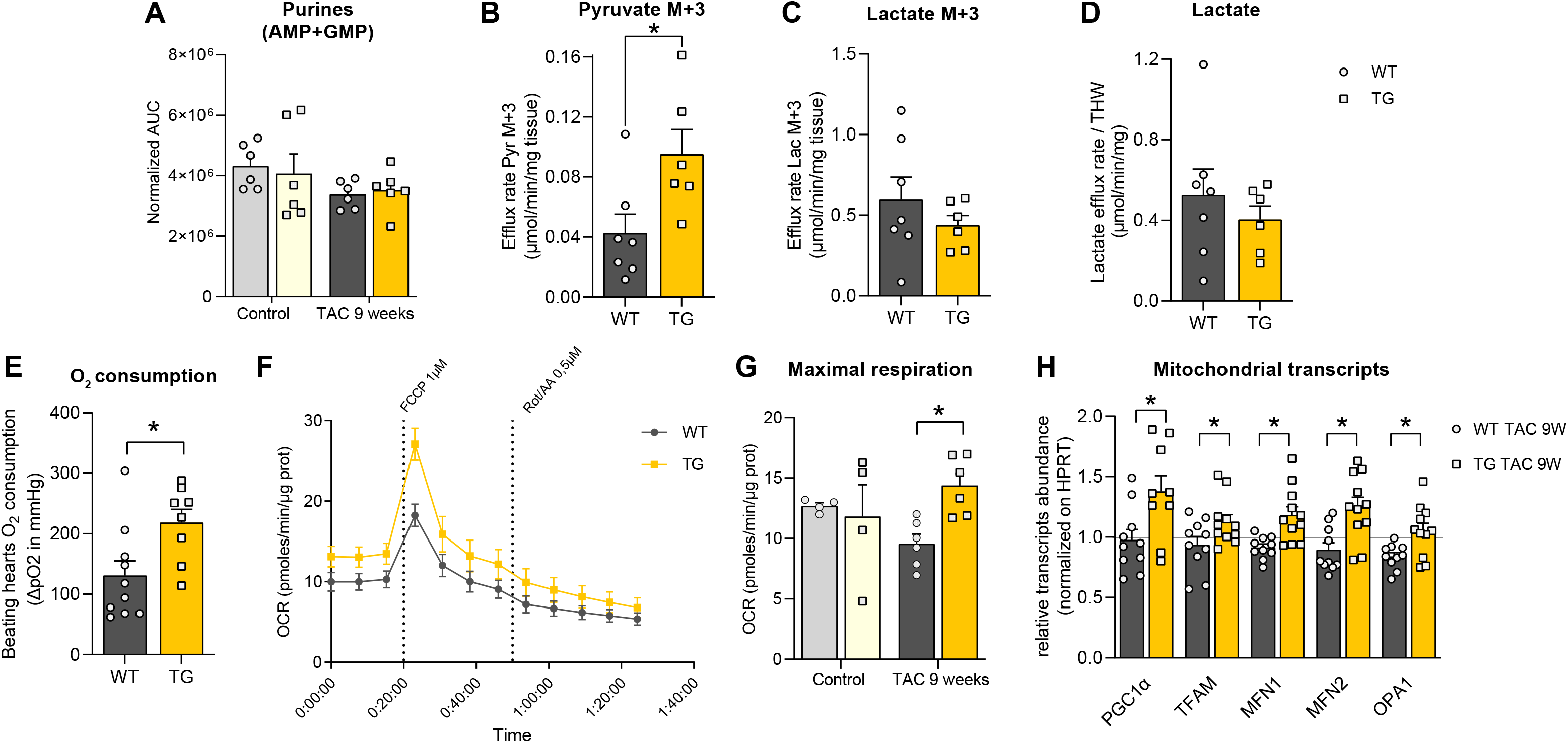
Cardiac β3AR promotes glucose oxidation and mitochondrial biogenesis. **– (A)** LC/MS determination of purines abundance in cardiac extracts from 9-week TAC or Control β3AR-Tg or wild-type mice. **(B&C)** Efflux rate of **(B)** pyruvate M+3 and **(C)** lactate M+3 and **(D)** lactate efflux rate in effluent of Langendorff-perfused hearts from β3AR-Tg and WT mice 9 weeks post-TAC perfused for 25 min with [U-^13^C_6_]-Glucose. **(E)** Oxygen consumed by Langendorff perfused hearts from 9-week TAC β3AR-Tg and WT mice **(F-G)** Assessment of mitochondrial respiration of AVM from 9-week TAC β3AR-Tg and WT mice in medium containing 11mM Glucose and 1ng/L Insulin by Seahorse analyzer; **(F)** Oxygen consumption rate (OCR); **(G)** maximal respiration **(H)** Transcripts abundance of markers of mitochondrial biogenesis (PGC1α, TFAM) and fusion (Mitofusin 1, Mitofusin 2, Opa 1) in AVM isolated from β3AR-Tg and WT mice 9 weeks post-TAC ; normalized levels (to HPRT) are reported as relative to levels in AVM from Control WT non-operated hearts (set as 1). Results are expressed as means ± SEM with *P < 0,05 calculated by Two-way Anova corrected for multiple comparisons with Sidak’s test, except for (C-F), analyzed by unpaired t-test.

### Cardiac β3AR transactivates the transcription factor NRF2 via NO signalling to exert its anti-hypertrophic function

RNAseq analysis of the transcriptome of AVM isolated from 9-week TAC β3AR-Tg and WT further revealed distinct expression patterns between the two genotypes (**Fig 7A**). Based on the 305 genes differentially regulated (**Fig 7B**), we performed an over-representation analysis (ORA) based on the TRRUST database ^23^ to identify putative transcription factors driving the differential expression. The results showed an overrepresentation of transcripts targeted by specific transcription factors, including SREBPF1, PPARlll and NRF2 (NFE2L2) among the top candidates (**Fig 7C**). Notably, NRF2 (*NFE2L2*), by binding to the canonical Antioxidant Response Element (ARE) is implicated in the regulation of key genes affecting glucose uptake, glycogenolysis, pentose phosphate pathway, ROS catabolism, mitochondrial function and biogenesis (**Fig 7D**). Therefore, we next examined the involvement of this transcription factor in the protection conferred by the cardiac β3AR.

**Figure 7.**
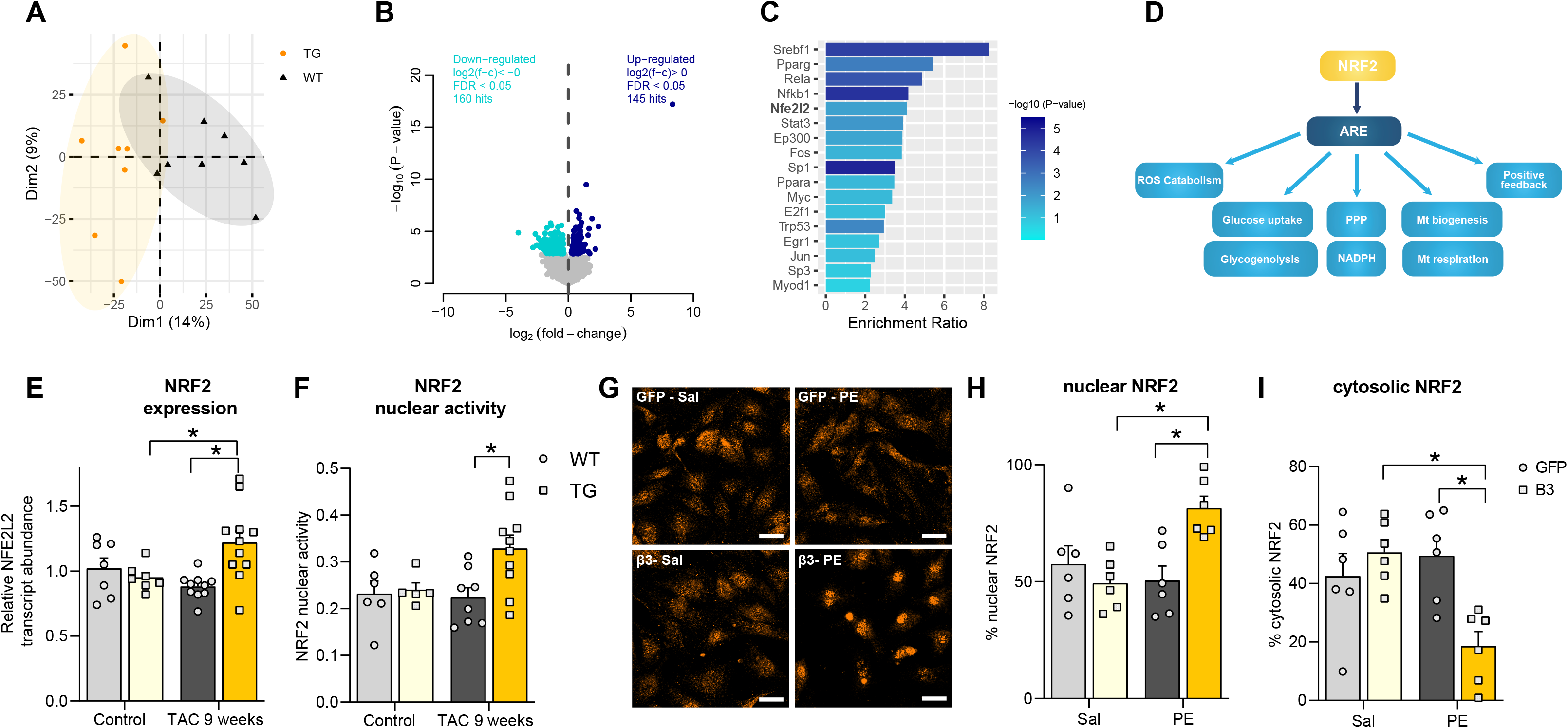
Cardiac β3AR activates nuclear translocation of the transcription factor NRF2. **– (A-C)** RNAseq of AVM isolated from β3AR-Tg and WT mice 9 weeks post TAC. **(A)** Principal Component Analysis (PCA) where biological replicates are marked in different colors according to subgroups: β3AR-Tg (orange) and WT (black) **(B)** Volcano plot of differentially expressed genes between β3AR-Tg and WT **(C)** Transcription factors enrichment ratio (β3AR-Tg vs. WT) computed with an over-representation analysis (ORA) of modulated genes by reference to the TRRUSTv2 database of transcriptional regulatory interactions ^23^. **(D)** Metabolic and intracellular pathways regulated by the transcription factor NRF2 (NFE2L2) **(E)** NRF2 (NFE2L2) transcripts abundance in AVM isolated from β3AR-Tg and WT mice 9 weeks post-TAC relative to levels in AVM from Control WT non-operated hearts (set as 1) **(F)** Binding capacity to antioxidant response element (ARE) measured in nuclear extracts from left ventricles of β3AR-Tg and WT as a proxy of NRF2 nuclear activity. **(G-I)** NRF2 Immunostaining in NRVM with adenoviral expression of human β3AR (Ad-β3AR) or control (Ad-GFP) **(G)** representative stainings of NRF2 (orange)(scale 20µm) **(H)** percentage of NRF2 nuclear localization **(I)** percentage of NRF2 cytosolic localization. Results are expressed as means ± SEM with *P < 0,05 calculated by Two-way Anova corrected for multiple comparisons with Sidak’s test, except for (C-F), analyzed by unpaired t-test.

NRF2 (*NFE2L2*) expression was upregulated in AVM from 9-week TAC β3AR-Tg hearts compared with WT (**Fig 7E**). NRF2 activity measured in nuclear extracts of left ventricles showed increased binding to specific antioxidant response element (ARE) regions in 9-week TAC β3AR-Tg hearts (**Fig 7F**). This was replicated in a model of neonatal rat ventricular myocytes (NRVM) combining adenoviral expression of human β3AR and treatment with the pro-hypertrophic agent, phenylephrine, with an enhanced translocation of NRF2 to the nucleus assessed by immunostaining (**Fig 7G-J**). Silencing of NRF2 with a siRNA (yielding more than 70% downregulation of NRF2) (**Fig 8A**) triggered a hypertrophic response in control cardiac myocytes with a magnitude similar to that of neurohormonal stress (**Fig 8B**). Notably, phenylephrine treatment of these cells failed to further enhance myocyte area suggesting that lower NRF2 activity recapitulates the hypertrophic response seen in neurohormonal stress. Next, the effect of silencing of NRF2 was examined in β3AR-expressing myocytes treated with phenylephrine. Loss of NRF2 abrogated the protective effect of β3AR with an enlargement of cardiac myocytes area similar to controls, thereby confirming the implication of NRF2 in β3AR anti-hypertrophic action (**Fig 8C**).

**Figure 8.**
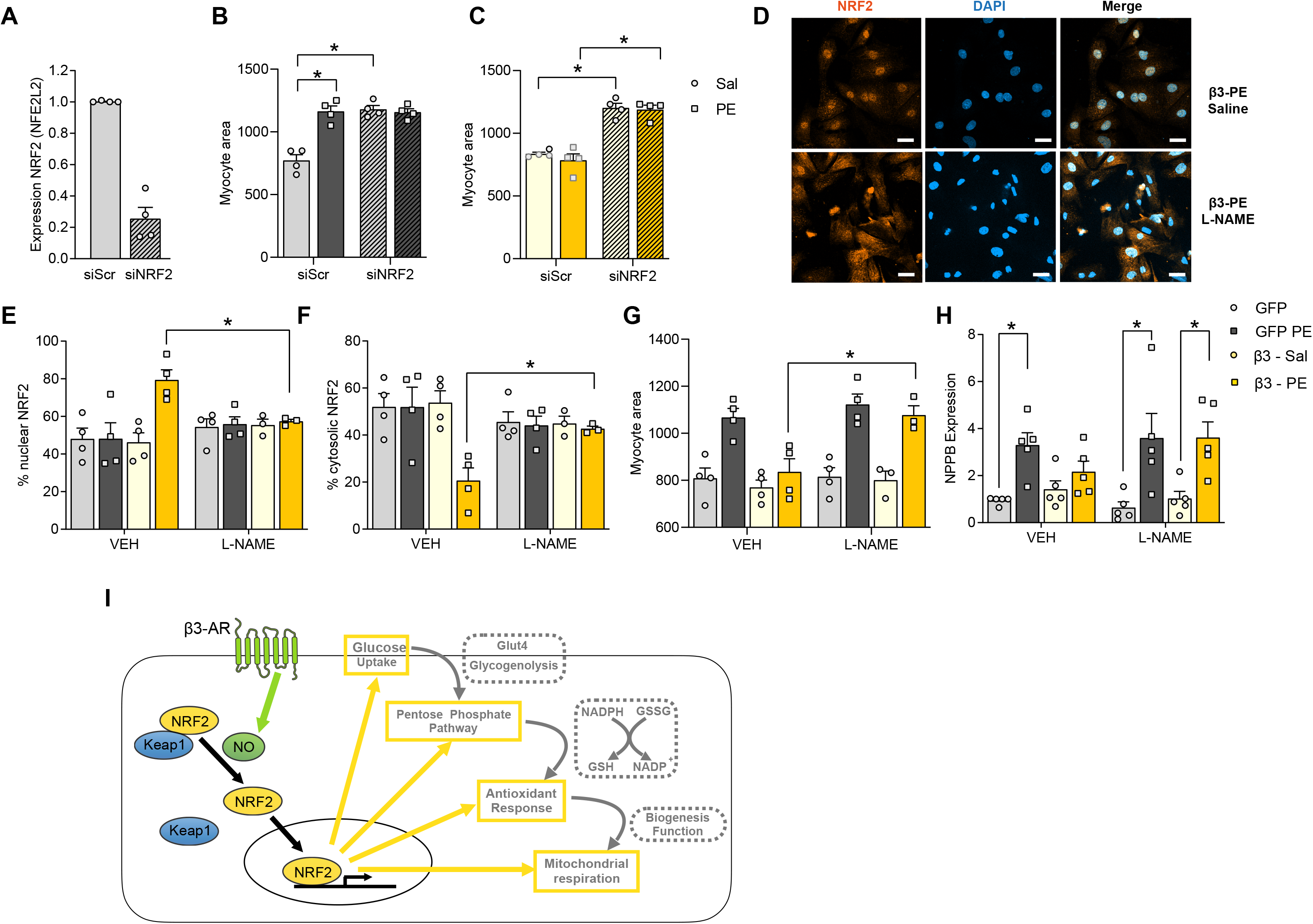
Cardiac β3AR regulates NRF2 translocation via NO signalling. **– (A)** NRF2 (NFE2L2) transcript levels upon treatment of NRVM with siRNA targeting NRF2 or scramble siRNA and **(B-C)** associated myocyte area upon treatment with PE (50µM) or Saline of **(B)** neonatal myocytes infected with Ad-GFP or **(C)** Ad-β3AR. **(D-G)** Effect of L-NAME (NOS inhibitor) administration (100µM) to NRVM adenovirally infected with Ad-β3AR or Ad-GFP and stimulated or not with phenylephrine; **(D)** representative staining of NRF2 (orange) and DAPI (blue) in PE-treated Ad-β3AR without (upper row) or with L-NAME treatment(lower row)(scale 20µm); **(E)** percentage of NRF2 nuclear localization; **(F)** percentage of NRF2 cytoplasmic localization; **(G)** associated myocyte area; and **(H)** Transcripts abundance of Nppb (BNP, as marker of hypertrophy) in NRVM infected with Ad-GFP or Ad-β3AR and treated with phenylephrine (PE ; 50µM) or Saline combined with L-NAME treatment (100µM) or vehicle (VEH). Results are expressed as means ± SEM with *P < 0,05 calculated by Two-way Anova corrected for multiple comparisons with Sidak’s test.

**Figure 9.** A NRF2/β3-adrenoreceptor axis drives a sustained antioxidant and metabolic rewiring through the pentose-phosphate pathway to alleviate cardiac stress.

As we and others previously demonstrated the coupling of β3AR to nitric oxide synthase (NOS) and nitric oxide (NO) production in cardiac myocytes, ^9, 10, 24^, and NO was shown to control nuclear translocation of NRF2 in other cell types ^25–29^, we tested the implication of NOS in the protection mediated by β3AR through NRF2 transactivation, by using the NOS inhibitor, L-Nitro-Arginine-Methyl-Ester (L-NAME), incubated on cardiac myocytes stimulated or not with phenylephrine. As shown in **Fig 8D-F**, the nuclear translocation of NRF2 in β3AR-expressing myocytes under neurohormonal stimulation was abrogated upon treatment with L-NAME, together with the loss of β3AR’s anti-hypertrophic effect assessed either by myocyte surface determination or hypertrophic marker (BNP) expression (**Fig 8G-H**). Altogether, this confirmed the implication of NOS enzymes in the translocation of NRF2 mediated by β3AR.

### Transcript abundance of elements of the PPP are enriched in human cardiac myocytes from heart failure patients who recover after left ventricular assist device implantation

Human heart failure entails profound myocardial remodeling associated with increased oxidant stress. Among therapeutic approaches, implantation of a left ventricular assist device is known to improve cardiac function and to produce “reverse remodeling”, including regression of cardiac hypertrophy ^30–33^. We then asked whether such beneficial effect is associated with changes in gene expression that would, to some extent, recapitulate our observations in β3AR-expressing hearts. We analyzed single-cardiac myocyte nucleus RNA sequencing data from a recently published dataset generated from patients with heart failure who recovered left ventricular systolic function after left ventricular assist device (LVAD) implantation. The output was compared with a similar dataset obtained in the same study from patients who did not recover LV function. The patients’ characteristics are detailed in ^22^. After computing differential gene expression, an over-representation analysis (ORA) was performed and the enrichment ratio of KEGG pathways plotted as illustrated in **Suppl Fig S5**. Transcripts associated with the Pentose Phosphate Pathway appeared among the top three differentially expressed gene sets, suggesting that regulation of the PPP parallels the recovery of LV function in LVAD responders (vs. non-responders). Among other significantly affected sets, genes associated with insulin resistance and cGMP-PKG signaling pathway (associated with nitric oxide production) are also differentially regulated, reminiscent of the improved insulin sensitivity associated with the NO-dependent activation of PPP in our model.

## Discussion

This work shows that the metabolism of glucose through the PPP can be recruited as a protective mechanism through a receptor-mediated pathway under cardiac stress. As such, it clarifies the beneficial role of the PPP and β3-AR against adverse myocardial remodeling and enhances the mechanistic understanding of its recruitment by signaling through nitric oxide and NRF2 (**Fig 8I**).

While previous studies have highlighted the advantage of activation of the oxidative part of the PPP for cell survival/function in yeast ^34^ or some mammalian cells ^35^, particularly cancer cells ^36^, its involvement in the regulation of oxidant stress or intermediate metabolism in whole tissues or organs in pathophysiological situations was less clear. As it readily produces NADPH, the PPP can both supply reductive equivalents to feed antioxidant mechanisms, but also provide the same NADPH to pro-oxidant NOX enzymes; in addition, the PPP provides intermediates (e.g. ribose-5P) for the synthesis of nucleotides, which can sustain cellular division and proliferation (e.g. in cancer cells)^37^, but also promote DNA and RNA synthesis that may promote hypertrophy in poorly dividing cardiac muscle cells ^38^. These diverging effects may, in part, explain the contradictory results observed previously in cardiac models with genetic deletion or inhibition of the key enzyme initiating the PPP, G6PD ^39, 40^. The current study clarifies these controversial issues in showing that activation of the β3AR pathway steers glucose metabolism towards beneficial effects under stress through antioxidant protection and preservation of mitochondrial respiration.

Enhanced myocardial glucose uptake is typically observed in the remodeling myocardium during the development of hypertrophy, ultimately culminating in heart failure ^41, 42^ Together with a shift away from fatty acid oxidation and increased reliance on glucose as metabolic fuel, the increases in glycolysis and in the branching hexosamine biosynthetic pathway are thought to contribute to hypertrophy, while the overall decreased oxidative metabolism impairs cardiac energetics ^43–45^. Contrary to reduced mitochondrial oxidation ^4, 5, 46^ the accrued influx of glucose mediated by β3AR appears both to fuel the pentose phosphate pathway and to increase mitochondrial respiration and was not associated with an increase in the hexosamine biosynthetic pathway. These alterations would improve cardiac energetics by readjusting the level of glucose oxidation, energetically the most efficient pathway with a high yield of ATP production for a lower O _2_ consumption than fatty acids oxidation ^47^ thereby improving cardiac efficiency ^48–50^. Increased uptake and oxidation of glucose can also promote the synthesis of aspartate from intermediates of the TCA; together with ribose-5P from the PPP, this was shown to increase *de novo* nucleotide synthesis and pro-hypertrophic effects of glucose utilization in cardiac myocytes treated with phenylephrine ^51^. This effect was fully antagonized by forced fatty acid oxidation upon genetic deletion of ACC2. Notably, these effects were independent of insulin ^51^. In contrast, here, β3AR activation of the PPP did not alter levels of nucleotides, but restored insulin sensitivity and glucose uptake under stress through Glut4 expression, suggesting a different route of glucose disposition. Also, we previously showed that under neurohormonal stress, cardiac β3AR sustains the activation of AMPK ^52^, a known inhibitor of ACC2, consistent with an anti-hypertrophic effect. Together, these effects may play a part in the protective effect of β3AR as insulin insensitivity has been shown to precede systolic dysfunction ^6, 53^ and reduced Glut4 has been associated with the development of cardiac hypertrophy ^54^.

Recruitment of the oxidative phase of the PPP would provide NADPH as required by intracellular antioxidant systems to reduce ROS, particularly H _2_O_2_ through glutathione peroxidase and thioredoxin ^55^. While NADPH can also be used by the NADPH oxidase (NOX) isoforms to generate ROS, our direct measurements of oxidant markers *ex vivo* and ROS in cardiac cells expressing β3AR clearly showed a decrease in intracellular oxidants under stress. Again, the sustained activation of AMPK observed in β3AR-expressing cells ^52^ may have preserved the redox balance since AMPK is known to negatively regulate NOX activity ^56, 57^.

Several additional features probably favored the recruitment of the PPP by β3AR under stress. A recent study highlighted the importance of the redox switch within GAPDH that supports the reversible inactivation of this key glycolytic enzyme by endogenous H _2_O_2_, necessary for derivation of glucose towards the PPP for antioxidant protection in cancer cells ^36^. Notably, GAPDH has long been known to be also sensitive to nitric oxide, that reversibly inactivates the enzyme by nitrosation of a key cysteine residue within its catalytic site ^58^. Together with an initial drop of NADPH/NADP+ ratio under stress, favoring G6PD dimerization and recruitment of reserve flux in the PPP ^59^, β3AR production of NO ^9–11^ may produce the necessary switch on GAPDH to allow glucose derivation to the PPP. Allowing this metabolic rewiring to be sustained under chronic stress, however, requires long-lasting effects. Our transcriptomic analysis in 9-week TAC β3AR myocytes indeed identifies upregulation of the expression of several key enzymes of the oxidative PPP that would sustain the increased flux through the pathway. Our over-representation analysis identified several putative transcription factors accounting for these expressional changes. Among the top candidates, SREBPf1, a transcription factor implicated in sterol metabolism, was previously shown to be part of the transcriptional program and anti-hypertrophic effect induced by activating Liver X receptor in cardiac myocytes and mice under chronic TAC ^60^. Peroxisome proliferator-activated receptor-gamma (PPARγ), a member of the nuclear receptor superfamily, is well known to be implicated in the regulation of mitochondrial biogenesis and metabolism through the PPAR/PGC1α complex, in addition to ancillary, non metabolic effects that have been implicated in antioxidant ^61^ and cardiac anti-hypertrophic effects ^62, 63^. Of particular interest is our identification of NRF2 from the ORA analysis of our transcriptomic data, which we further validated functionally in isolated cardiac myocytes.

Nuclear factor(erythroid-derived2)-like2 NRF2 is a well described transcriptional regulator of enzymes of the pentose phosphate pathway and antioxidant response ^64^. More recently, the NRF2/ARE pathway was reported to regulate mitochondrial biogenesis through NRF1 regulation ^65^ and PGC1α ^66^ as well as glycogenolysis ^67^. These effects are consistent with our observation of decreased glycogen content, together with upregulation of glycogenolytic enzymes and mitochondrial respiration. Loss-of-function experiments in NRF2 ^-/-^ mice have yielded contrasting results in the unstressed heart ^68–70^, but an anti-hypertrophic action of NRF2 in cardiac myocytes was previously reported under hemodynamic ^69, 71^ and neurohormonal stress ^70, 72^. Maladaptive remodeling mediated by NRF2 has occasionally been reported. Further investigations revealed that adverse remodeling is restricted to the specific context of impaired autophagy ^73–75^ which, however, is preserved under β3-AR expression ^52^. In isolated cardiac myocytes, silencing of NRF2 consistently leads to a marked hypertrophic response ^70^, as observed in the present study. However, the mechanistic connection to changes in the intermediate metabolism was unknown. Therefore, ours is the first evidence for a receptor-mediated activation of NRF2 that orchestrates an antioxidant response and metabolic rewiring resulting in preserved mitochondrial respiration and protection from cardiac hypertrophy under stress.

Mechanistically, our study demonstrates that NRF2 nuclear translocation is dependent on nitric oxide synthase activity. Coupling of β3AR with Gαi and downstream NOS (eNOS and nNOS) in cardiac myocytes has been well established ^9–11, 76^. Accordingly, our present data using the NOS inhibitor L-NAME confirm that β3AR regulation of NRF2 translocation results from an increased nitric oxide (NO) production. NO-mediated NRF2 activation has been identified in other cell types ^25–29^, in which S-nitrosation (SNO) of a redox-sensitive cysteine residue in Keap1, the dominant repressor of NRF2 ^77, 78^ has been suggested to facilitate its dissociation from NRF2 thereby allowing its nuclear accumulation.

Of note, the attenuated hypertrophic response in β3AR-expressing hearts did not compromise cardiac function, as reflected by the maintained ejection fraction and fractional shortening under stress, probably owing to improved metabolic efficiency, as demonstrated above. Together with our analysis of human single cell data, previous observations of increased expression of enzymes of the PPP and reduced ROS levels in the myocardium of patients who favorably responded to LVAD implantation ^79^suggestthatsimilarmechanismsmaybeatplayinhuman disease. Indeed, β3AR are upregulated in patients with heart disease ^17^ and recent clinical trials showed that a 6-month treatment with mirabegron, a specific β3AR agonist, improves ejection fraction in heart failure patients with NYHA class III–IV and severe LV dysfunction at baseline (LVEF < 40%) ^19^. In addition, 1-week treatment was sufficient to significantly improve cardiac index and vascular pulmonary resistance in patients with LVEF < 35% ^20^.

In conclusion, moderate expression of β3AR in cardiac myocytes activates a coordinated antioxidant response and metabolic rewiring through NRF2 to preserve myocardial energetics and functional integrity in the face of hemodynamic stress.

## Supplemental methods

### Animals

All protocols were carried out in accordance with the Guide for the Care and Use of Laboratory Animals published by the U.S. National Institutes of Health (NIH) and the European Directive 2010/63/EU and were approved by local ethical committees. Heterozygous male β3AR-Tg mice harboring a human ADRB3 transgene under the control of the alpha-myosin heavy chain (Myh6) promoter (previously described ^11, 80^ and wild-type (WT) littermate controls were obtained from Dr D. Langin, Univ. Toulouse (FR). Ascending Transverse aortic constriction (TAC) was performed on age-matched 12-16 week-old male mice from both genotypes as described ^11^ and compared to control non-operated littermates. After anesthetizing the mouse with ketamine/xylazine, a constrictive band was placed and tightened around the aorta to the diameter of a 27 G needle. Animals were treated with 0,1mg/kg of buprenorphine after the surgery. Doppler measurements of trans-stenotic gradients were systematically performed at day 3, then week 3 and 9 post surgery. Only mice with a velocity between 2,5 and 4,5 m/s were kept into an experiment.

### Echocardiography

Two-dimensional echocardiography was performed wit h a high-resolution ultrasound system (Vevo 3100 VisualSonics; 40 Mhz probe) in anesthetized animals (1-3 % isoflurane) placed in supine position and fixed on limb electrodes to monitor heart rate (HR). LV parasternal long axis view was recorded in B-mode to measure LV internal volumes at end of diastole (LVEDv) and systole (LVESv), used to calculate the ejection fraction (EF; Bmode). M-mode view was performed to measure LV posterior wall (PW) and interventricular septum (IVS) thicknesses, LV internal dimensions (LVID), allowing the assessment of the fractional shortening (FS). LV mass was calculated as 1,053*[(LVIDd + PWd + IVSd)³-LVIDd³].

### Histomorphometry

Heart sections were fixed overnight in 4 % formaldehyde, embedded in paraffin. Myocardial samples were serially cut in 5 µm thick sections and stained with rhodamine-conjugated–wheat germ agglutinin (WGA; Vector RL-1022; vector labs) according to standard protocol. Stained sections were digitized with a slide scanner Axioscan. For each heart, area of 200 cells oriented transversally were measured on three sections distant from 100 µm, using Visiopharm software.

### Adult ventricular myocyte isolation

Adult mice were anesthetized with ketamine/xylazine. Hearts were quickly excised, cannulated and mounted onto a Langendorff system. Subsequently, hearts were perfused at 37 °C with a Liberase-containing solution (113 mM NaCl, 4,7 mM KCl, 0,6 mM KH _2_PO_4_, 0,6 mM Na _2_HPO_4_, 1,2 mM MgSO _4_, 12 mM NaHCO _3_, 10 mM KHCO _3_, 30 mM Taurine, 25 µg/ml Liberase TH (Roche), 10 mM BDM, 5,5 mM Glucose and 10 mM HEPES/NaOH, pH 7,46) at a constant rate of 2,5 ml/min. Once digested, cells were mechanically separated and filtered. Thereafter, cytosolic Ca ^2+^ levels were gradually restored. Viable rod-shaped cardiac myocytes were separated by sedimentation and plated on laminin-coated wells in M199 medium containing 10 mM BDM, 5% FBS and Penicillin/ Streptomycin. After one-hour plating, medium was replaced by culture medium (M199 medium, 10 mM BDM, 1X Insulin-transferrin-selenium, ITS, 1 mg/mL BSA and 10 mM HEPES) before being processed for further experiments.

### In vivo myocardial ^18^FDG and ^18^FTO uptake measurements

In vivo cardiac glucose and fatty acid uptake were assessed using whole body positron emission tomography (PET) imaging performed on a dedicated small-animal PET scanner (Mosaic; Philips Medical Systems) with a spatial resolution of 2,7 mm (FWHM)^81^. Animals fed ad-libitum received a dose of 10–15 MBq/animal (Betaplus Pharma) of 2-deoxy-2-[^18^F]-d-glucose (^18^F-FDG) or 6–12 MBq/animal (Betaplus Pharma) of 18-^18^F-Fluoro-4-Thia-Oleate (^18^F-FTO) ^82^ by tail-vein injection. Animals were kept under a heating lamp and on a heated pad before acquisition. Thereafter, acquisition was performed under continuous 2% isoflurane anesthesia on a heated animal bed. A 10-min emission scan was performed 1 h after tracer injection for ^18^F-FDG and a 30-min emission scan was performed 90 min after ^18^F-FTO. Anesthetized mice were then transferred on the same bed to a microCT (NanoSPECT/CT Small Animal Imager, Bioscan, Inc.) for anatomic reference of PET images. PET images were reconstructed using a fully three-dimensional interactive algorithm (3D-RAMLA) in a 128 x 128 x 120 matrix, with a voxel of 1mm3. Cardiac tracer uptake was assessed on the reconstructed images and expressed as standardized uptake value (SUV) max calculated as cardiac activity/(injected dose/bodyweight)). SUVs were computed from a volume of interest automatically drawn around the whole heart using a 60%-threshold of the maximum, manually adjusted if necessary (MIM 6.7 - MIM Software Inc., Cleveland, OH).

### ^18^FTO synthesis and validation

#### Synthesis of methyl 3-(hex-5-en-1-ylthio)propanoate (1)

To a mixture of acetone (30 mL) and DMSO (10 mL) was added sequentially K _2_CO_3_ (2,8 g, 20,26 mmol, 2.4 eq), 6-bromohex-1-ene (1,6 g, 10 mmol, 1.2 eq) and methyl 3-mercaptopropanoate (1 g, 8,32 mmol, 1 eq). The resulting mixture was stirred for 56h at room temperature, then filtered on Celite®, and washed with acetone (2x 10 mL). The mixture was concentrated under vacuum, then DCM (80 mL) was added and the solution was washed with water (4x 20 mL). The organic layer was dried over MgSO _4_, and filtered. The volatiles were removed under vacuum. The residue was purified by silica gel chromatography (PE:EtOAC, 100:0 to 80:20) to afford the pure product (1,4 g, 85%)

#### Synthesis of methyl (Z)-3-((14-bromotetradec-5-en-1-yl)thio)propanoate (2)

In a glovebox, a 20 mL vial was charged with (1) (0,200 mg, 1 mmol), 10-bromodec-1-ene (0,331 mg, 1,5 mmol), and THF (10 mL). Then Hoveyda-Grubbs catalyst® M2001 (6,5 mg, 0,01 mmol, 1 mol %) was added and the reaction was stirred at 35 °C in an open vial for 5 hours. The vial was removed from the glovebox and the solvent was removed in vacuo. The residue was purified by silica gel chromatography (PE:EtOAC, 100:0 to 85:15) to afford a mixture of the product and the 10-bromodec-1-ene starting material. The mixture was purified by reverse-phase chromatography using a C18 column (H2O:ACN, 10:90 to 0:100) to afford the product (0,165 mg, 40%).

#### Radiofluorination

The optimal bromoester precursor (2) was then processed for radiofluorination according to already described method ^82^. Briefly, cyclotron-produced ^18^F-fluoride was dried down with Kryptofix 2.2.2, acetonitrile, and K _2_CO_3_ solution in H _2_O. A solution of (2) in acetonitrile was added and heated at 75°C for 15 min. After brief cooling, subsequent hydrolysis was performed by 0,2N KOH addition and heating it at 90°C for 4 min. The mixture was cooled, acidified with concentrated acetic acid, filtered, and applied to the semipreparative HPLC column (C-18). The 18F-fluoro–fatty acid fraction was diluted in water and trapped on a C-18 Sep-Pak cartridge. After washing the product was eluted from the Sep-Pak in ethanol, then briefly evaporated to reduce EtOH content before ^18^F-FTO formulation in 1% albumin in isotonic NaCl solution and filtered through 0,22-µm filter.

#### ^18^F-FTO validation

Control animals were subjected to intraperitoneal injection of Etomoxir 30mg/kg body weight or saline 2 hours before ^18^F-FTO injection leading to a decrease > 50% in cardiac SUV.

### 2-^3^H Glucose uptake in AVM

Glucose uptake was measured, as described previously ^83^, by following the [2-^3^H] glucose (Perkin Elmer) detritiation rate through the detection of tritium labelled-H _2_O formed during rapid isomerization by the action of phosphohexose isomerase ^84, 85^. Briefly, for the last 30 min of treatment, 0,2 µCi/ml of [2-^3^H] glucose was added to the culture medium containing 5,5 mM of glucose with or without 50nM Insulin. Tritiated water present in the supernatant was separated from tritiated glucose using anion exchange chromatography using anion exchange resin placed in suspension in borate buffer beforehand and then measured by a scintillation counter. Results were normalized on input and protein quantitation.

### Neonatal rat ventricular myocytes isolation and associated procedures

#### Neonatal rat ventricular myocytes isolation

were isolated from 10-15 hearts of neonatal pups following a protocol adapted from ^86^. Heart ventricles were removed from neonatal pups, minced into 1-mm ^3^ pieces, and subjected to overnight digestion with collagenase type II (Worthington Bioscences) and trypsin (Gibco). Cells were separated by using Percoll gradient centrifugation and selective attachment of fibroblasts. Myocytes were cultured in DMEM supplemented with 10% fetal bovine serum (FBS), 1% of penicillin/ streptomycin and supplemented for the first 24 hours with 10 μM AraC to prevent fibroblast and endothelial cells proliferation. Cells were maintained in a humidified atmosphere of 5% CO _2_ at 37°C.

#### NRF2/NFE2L2 silencing

Approximately 8h post-isolation, myocytes were transfected with siRNA targeting NRF2 (sc-156128, SantaCruz) using Lipofectamine RNAimax reagent (Thermofisher). In 6 well-plates containing 800 000 myocytes, transfection with 25pmol, 50pmol and 75pmol of siRNA led to a similar downregulation of the target of about 70-80%. Therefore, 25pmol was used in the subsequent experiments for siRNA NRF2 and scramble siRNA (sc-37007, Santacruz).

#### Adenoviral expression of human β3AR

Approximately 20h post-isolation, myocytes were transferred to serum-free media and infected with a recombinant adenovirus containing a bi-cistronic construct coding the human ADRB3 cDNA construct and GFP ^10, 11^ at a multiplicity of infection (MOI) of 1,5 plaque forming units (PFU) per cell; an adenovirus encoding only GFP was used as control in all experiments.

#### Treatments

Approximately 24h after infection, myocytes were treated with either phenylephrine-PE (50μM), L-Name (100µM) or saline in freshly added serum-free media.

### 1-^14^C-Palmitate uptake in AVM

Freshly isolated AVM were resuspended in 10mL of medium 3 in MKR solution (113 mM NaCl, 2,6 mM KCl, 1,2 mM KH _2_PO_4_, 1,2 mM MgSO _4_, 10 mM NaHCO _3_, 0,8 mM CaCl _2_, 10 mM HEPES/NaOH, pH 7,46 and 5,5mM Glucose). For each condition, 200 000 cells are incubated with MKR in fatty acid free glass tubes for which at t=0 MKR was supplemented with 20µM palmitate complexed with fatty acid free BSA (Ratio FA:BSA, 1:3,5) and containing 0,03 µCi of 1-^14^C-Palmitate (Perkin-Elmer) prepared as previously described ^87^. In order to avoid loss of ^14^C labeling through fatty acids oxidation, palmitate uptake was stopped after 5 minutes by the addition of MKR medium supplemented with phloretin 0,2mM. Cells were thoroughly washed, pelleted and lyzed before ^14^C measurement by a scintillation counter. Results were normalized on input and protein quantitation. In order to ensure the viability of cellular preparation (membrane disruption) conditions with stimulated or inhibited uptake were included for each cellular preparation, by a 15-min pre-treatment with either Insulin 50nM for stimulation or Phloretin 400µM combined with SSO 400µM for inhibition.

### Glycogen content determination

Glycogen content was determined in left ventricular samples using the glycogen Assay Kit (ab65620, abcam). Shortly, glycogen was first extracted in 30% KOH followed by 2 hours incubation at 100°C. Glycogen was then precipitated by 2 volumes of 95% ethanol and then dissolved in H_2_O after centrifugation (10 min – 18 000xg – 4°C), a procedure repeated 2 times. Hydrolysis of glycogen was performed with glucoamylase to extract glucose. Thereafter, glucose was specifically oxidized to react with OxiRed probe for which absorbance was measured by spectrophotometer at 570 nm. Standard curve was used to quantify glycogen in the LV extracts.

### Glucose and insulin tolerance test

Mice underwent GTT and ITT after a fasting period of six and four hours, respectively. After initial glycemic measure by tail blood sampling, a bolus of glucose (2g/kg) or insulin (0,5U/kg) was injected into the intraperitoneal cavity. Thereafter, blood was sampled at 0, 15, 30, 60, and 120 min for glycemia using test strips with the associated Contour Next glucose meters (Ascensia). To avoid variability due to diurnal rhythms, measurements were consistently performed in the afternoon. Data are expressed as plasma glucose(mg/dl) for GTT and as % of basal glycemia for ITT.

### Serum analysis

#### Insulinemia

Blood was collected during GTT in animals 9 weeks post-TAC 30 minutes after glucose bolus injection and spun to isolate serum, then stored at − 80 °C. Serum samples were subsequently analysed for insulin levels using the Ultra sensitive mouse insulin ELISA kit (90080; Crystal Chem).

#### BNP measurements

Blood was collected during terminal procedures 9 weeks post TAC and spun to isolate serum, then stored at − 80 °C. Then, BNP levels were measured in the serum samples using the mouse BNP EIA kit (EIAM-BNP; RayBiotech).

### Ex vivo heart perfusion

Adult mice were anesthetized with ketamine/xylazine. Hearts were quickly excised while immersed in cold saline and a cannula was inserted into the aorta. Thereafter, hearts were perfused in the Langendorff mode while protected by an organ bath to maintain a physiological temperature of 37,5 °C. Hearts were perfused at a constant flow of 2,5mL/min of with a modified Krebs–Henseleit buffer containing: 110 mM NaCl, 4,7 mM KCl, 2 mM CaCl _2_ (in order to obtain [free Ca ^2+^] of 1,5 mM, 0,24 mM KH _2_PO_4_, 0,48 mM K_2_HPO_4_, 0,48 mM Na _2_HPO_4_, 1,2 mM MgSO _4_, 25mM NaHCO _3_ and 0,1 mM EDTA supplemented with 2 nM insulin, 0,2 mM pyruvate, 1,5 mM lactate, 0,5 mM lactate and 50 μM carnitine supplemented with 0,4mM palmitate complexed with BSA (4:1) and glucose using first 11mM unlabeled ^12^C-glucose then switched to 11 mM [U-^13^C_6_]-Glucose (Perkin-Elmer) during perfusion with labeled buffer as previously described ^88^. After a 5-min initial stabilization with unlabeled metabolic substrates during which the hearts spontaneously resume beating, the perfusion solution was switched to the ^13^C-labeled glucose for a duration of 5, 10 or 25 minutes in order to follow the kinetics of incorporation until reaching isotopic equilibrium. During the experiment, buffers were continuously pumped through an oxygenator connected to a gas cylinder (95% O _2_–5% CO_2_) to maintain perfusion buffers at pH 7,4 and a pO _2_ >500 mmHg. Oxygen consumed by the hearts perfused in the Langendorff mode was measured by subtracting pO _2_ measured in effluent (downstream the perfused heart) from pO _2_ measured in perfusate (upstream) using iSTAT blood analyzer (Abbott) with the corresponding CG8 ^+^ cartridges for blood gases assessment. At end point, effluents were collected and hearts were freeze-clamped with metal tongs and stored at - 80°C for subsequent metabolic measurements.

### [Lactate] determination

Effluent and influent perfusates were collected from the *ex vivo* hearts perfused in the Langendorff mode for 25 min and lactate standards were measured in Tris-hydrazine buffer (0,1M Tris; 10mM MgSO _4_; 5mM EDTA; 1M hydrazine hydrate; pH 8,5) complemented with 1,4mg/mL NAD (10621650001; Roche) and 0,15mg/mL of lactate dehydrogenase (10127884001; Roche). Enzymatic production of NADH,H^+^ was assessed by spectrophotometer absorbance reading at 340nm. Influent was subtracted from effluent [Lactate] to determine [Lactate] efflux by minute, knowing that the heart is perfused at a constant flow rate of 2,5mL/min, and normalized on heart weight measured before perfusion.

### Metabolic flux analysis and MPE determination

#### MPE G6P/F6P/PPP determination

Extraction and LC/MS have been performed based on the validated method to measure HBP described in detail ^89^ and modified to include the pentose phosphate pathway intermediates. Briefly, 50 mg of freeze-clamped heart tissue pulverized in liquid N _2_ were deproteinized with a mix of 1 ml of MeOH 70%. Samples included WT hearts (25 min, n=7; 10 min, n=4; 5 min, n=4) and TG hearts (25 min, n=6; 10 min, n=3; 5 min, n=3). After centrifugation (12,000g - 10 min - 4°C), the liquid phase was collected, and the pellet was extracted a second time in 450 µl of MeOH 70%. After vortex and centrifugation, the two liquid phases were combined and filtered through a 13-mm GD/X 0,45-µm syringe filter (Whatman, GE Healthcare), and filters were washed with 1 ml of MeOH 70%. The combined MeOH phases were evaporated under nitrogen until 200µl and kept at + 4°C overnight. Next, samples were centrifuged (3000g - 5 min), and a volume of 2 µL of the upper liquid phase was injected into the LC-MS (1290 Infinity HPLC equipped with a SeQuant ZIC-pHILIC PEEK-coated column (2,1x150 mm, 5 µm; EMD Millipore, Billerica, MA) and coupled with a 6530 accurate mass quadrupole-TOF (LC-QToF, Agilent Technologies) with a Dual Agilent Jet Stream ESI source. Chromatographic conditions and LC-MS operating conditions as well as mass correction were applied as previously described and has been validated for linearity, limit of detection, and intra- and interday reproducibility ^89^. MS signals were extracted using Mass Hunter Quantitative Analysis version B.07 from Agilent. ^13^C-Labeling of RR5P (Ribose/ribulose -5-phosphate), G6P (Glucose-6-phosphate), F6P (Fructose-6-phosphate), S7P (Sedoheptulose-7-phosphate) and UDP-GlcNAc were quantified by extracting the MS signals corresponding to the M and labelled M+ (number of ^13^C) isotopomers ions and expressed as molar percent enrichment (MPE). The m/z for the various metabolites are as follows: RR5P, 229,0119 (unlabeled) and 234,0284 (M+5); G6P, 259,0224 (unlabeled) and 265,0422 (M+6); F6P, 259,0224 (unlabeled) and 265,0422 (M+6); S7P, 289,0330 (unlabeled) and 296,0561 (M+7); UDP-GlcNAc, 606,0743 (unlabeled) and 612,0944 (M+6)

#### MPE Pyruvate/Lactate from effluent

Influent and effluent perfusates were used to assess ^13^C-enrichment in lactate and pyruvate arising from cytosolic glycolysis of exogenous [U-^13^C_6_]-glucose using GC-MS (Agilent 6890N GC coupled to a 5973N MS). Perfusates were treated with NaBD _4_ to reduce unlabeled (M) and [U-^13^C_3_] labeled (M+3) pyruvate into corresponding [^2^H] lactate (M+1) and [^2^H, U-^13^C_3_] lactate (M+4), respectively. The molar percent enrichment of the four different mass isotopomers of lactate were then combined with the [lactate] determination described above to determine pyruvate (M+3) and lactate (M+3) efflux after normalizing on heart weight and on perfusion rate (constant flow rate of 2,5mL/min) as described in details in the previously published method ^88^.

### Metabolomics

#### Unbiased metabolomics

Sample preparation: Metabolites from serum samples were extracted in microcentrifuge tubes after the addition of 3 volumes of 100% acetonitrile containing 3 internal standards [caffeine-(trimethyl-d9), succinic acid-2,2,3,3-d4 and N-acetyl-aspartic acid-2,3,3-d3] on one volume of serum. The samples were thoroughly mixed, sonicated and incubated at -20°C for 30 min, then centrifuged at 10,000g for 10 minutes at 4°C. The upper-phase was collected into a new microcentrifuge tube and 3 volumes of pure acetonitrile were added to the initial pellet for a second similar extraction. The combined upper-phases resulting from the two extractions were mixed and divided into four equal parts. Each part was dried down under a gentle stream of nitrogen at 30°C and kept frozen until analysis. Prior to analysis, the samples were reconstituted in [water:acetonitrile, 50:50 (v:v), 0,1% formic acid], [water:acetonitrile, 50:50 (v:v)], [water:acetonitrile, 25:75 (v:v), 0,1% formic acid] and [water:acetonitrile, 25:75 (v:v)] depending of the method used, “M1 POS”, “M1 NEG”, “M2 POS” and “M2 NEG”, respectively. The tubes were centrifuged at 10 000g for 5 minutes at 4°C to remove any particulates before being transferred into polypropylene autosampler vials. Liquid chromatography and mass spectrometry (MS-qTOF) analysis: 3µL of each reconstituted sample were injected and subjected to reverse phase (“M1”) or HILIC (“M2”) liquid chromatography using an UPLC ACQUITY Premier system (Waters) in both positive (“POS”) and negative (“NEG”) ionization modes. The separation columns, operated at 40°C, were an Acquity Premier HSS T3 column 1,8 µm, 2,1 x 100 mm (Waters) and an Acquity Premier BEH Amide 1,8 µm, 2,1 x 100 mm (Waters), for reverse phase and HILIC chromatography, respectively. Mass spectrometry data was acquired in profile (continuum) format over the mass range of m/z 50-1200 using a Waters Synapt XS high resolution Q-TOF mass spectrometer set in MSE resolution mode (scan time 0,1 sec, the low and high trap collision energy were 4V and a ramp between 20 and 50 V, respectively). A dual electrospray ionization (ESI) source was used in positive or negative mode. Capillary and sampling cone, voltages were set to 1 kV and 30 V or 1 kV and 25 V in positive and negative mode, respectively. Source temperature was set to 150 °C and desolvation temperature to 600 °C (550°C in negative ESI). Gas flow rates were set at 1200 L/h for the desolvation gas (1100 L/h in negative ESI) and 50 L/h for the cone gas. Acquisition of leucine enkephalin infused at 10ul/min through a lockspray probe allowed a real time mass correction (30 sec scan intervals). Each analytical batch contained blanks, quality controls (a reference standard mix and a pool of plasma samples used to monitor the performance of the analytical platform), study samples and samples obtained after pooling all the study samples (used for column conditioning and chromatogram alignment during data processing). Liquid chromatography was performed using a binary mobile phase system. The mobile phase A used for reverse phase liquid chromatography in positive electrospray (“M1 POS”) consisted of water with 0,1 % formic acid and the “M1 POS” mobile phase B consisted of acetonitrile with 0,1% formic acid. The composition of mobile phases used for the reverse phase chromatography operated in negative electrospray (“M1 NEG”) were the same except that formic acid was replaced by 0,1% acetic acid. The reverse phase separations (“M1”) in positive and negative electrospray were performed at 0,5 mL/min with an isocratic period of 1 min at 99% A, followed by three consecutive linear gradients: from 1% B to 15 % B over 2 min, from 15% B to 50 % B over 3 min and from 50% B to 95 % B over 3 min. The final composition of 95 % B was held constant for 4 min followed by a return to 99% A, kept for 1 min. The mobile phases used for the HILIC method (“M2”) in positive electrospray (“M2 POS”) consisted of [water:acetonitrile, 93:7 (v:v), 10 mM Ammonium Formate, pH 3,0] and [water:acetonitrile, 7:93 (v:v), 10 mM Ammonium Formate, pH 3,0] for mobile phases A and B, respectively. The composition used for the reverse phase chromatography operated in negative electrospray (“M1 NEG”) were the same except that the pH was adjusted to 9,0. The HILIC separations in positive and negative electrospray were performed at 0,7 mL/min with an isocratic period of 0,2 min at 100% B followed by two consecutive linear gradients, from 100% B to 80 % B over 8,3 min, from 80% B to 60 % B over 1 min and kept for 0,5 min. Then a return to 100% B, kept for 3 min of equilibration. Post-analytical processing: Raw LC-MS data were imported and processed in Progenesis QI software (Nonlinear Dynamics), which performed chromatogram alignment, peak picking, ion deconvolution, normalization, peak annotation and statistical analysis. Thresholds of area under the curve > 1000 were set and yielded the following number of detected metabolites: for Control WT vs Control TG (M1 neg : 1886 ; M1 pos : 3881 ; M2 neg : 785 ; M2 pos : 2411) TAC 9W WT vs Control TG (M1 neg : 2022 ; M1 pos : 3690 ; M2 neg : 905 ; M2 pos : 2405) and Control WT vs TAC WT (M1 neg : 2165 ; M1 pos : 4068 ; M2 neg : 944 ; M2 pos : 2448). Volcano plot and FDR threshold of significance were performed on Graphpad Prism.

#### Targeted metabolomics by Ion pairing

was performed on complete freeze-clamped hearts pulverized in liquid N_2_ for which metabolites were extracted. Subsequently to assess metabolic pathways using glucose as a substrate 10 μl of each sample was loaded into a Dionex UltiMate 3000 LC System (Thermo Scientific Bremen,Germany) equipped with a C-18 column (Acquity UPLC -HSS T3 1. 8 μm; 2,1 x 150 mm, Waters) coupled to a Q Exactive Orbitrap mass spectrometer (Thermo Scientific) operating in negative ion mode as previously published ^90^. A step gradient was carried out using solvent A (10 mM TBA and 15 mM acetic acid) and solvent B (100% methanol). The gradient started with 5% of solvent B and 95% solvent A and remained at 5% B until 2 min post injection. A linear gradient to 37% B was carried out until 7 min and increased to 41% until 14 min. Between 14 and 26 minutes the gradient increased to 95% of B and remained at 95% B for 4 minutes. At 30 min the gradient returned to 5% B. The chromatography was stopped at 40 min. The flow was kept constant at 0,25 mL/min and the column was placed at 40°C throughout the analysis. The MS operated in full scan mode (m/z range: [70.0000-1050.0000]) using a spray voltage of 4,80 kV, capillary temperature of 300°C, sheath gas at 40.0, auxiliary gas at 10.0. The AGC target was set at 3.0E+006 using a resolution of 140000, with a maximum IT fill time of 512 ms.

To assess specifically NADPH and NADP+, mass Spectrometry measurements were performed using Dionex UltiMate 3000 LC System (Thermo Scientific) coupled to a Q Exactive Orbitrap mass spectrometer (Thermo Scientific) operated in negative mode as previously described ^91, 92^. 10 μl sample was injected onto a Poroshell 120 HILIC-Z PEEK Column (Agilent InfinityLab). A linear gradient was carried out starting with 90% solvent A (acetonitrile) and 10% solvent B (10 mM Na-acetate in mqH2O, pH 9.3). From 2 to 12 min the gradient changed to 60% B. The gradient was kept on 60% B for 3 minutes and followed by a decrease to 10% B. The chromatography was stopped at 25 min. The flow was kept constant at 0.25 ml/min. The columns temperature was kept constant at 25 degrees Celsius. The mass spectrometer operated in full scan (range [70.0000-1050.0000]) and negative mode using a spray voltage of 2.8 kV, capillary temperature of 320°C, sheath gas at 45, auxiliary gas at 10. AGC target was set at 3.0E+006 using a resolution of 70000. Data collection was performed using the Xcalibur software (Thermo Scientific). For both Ion pairing and HILIC, the data analyses were performed by integrating the peak areas (El-Maven – Polly - Elucidata) according to the standard procedures of the VIB Metabolomics Core - Leuven. Subsequently, peak areas named areas under the curve (AUC) were corrected for heart weight and either plotted or used to calculate z-score (as population mean value subtracted from the individual raw value and divided by the standard deviation). Retention times and m/z of all metabolites are reported in Suppl table 2.

### RNA extraction and real-time reverse transcription–polymerase chain reaction

Total RNA was extracted from AVM using TRI-Reagent (Fermentas, Alost, BE) followed by PureLink RNA micro Kit (12183018A; Invitrogen) and from NRVM using Maxwell RSC simplyRNA Cells Kit (AS1390; Promega) with both kits including a DNAse treatment. Eight hundred ng of total RNA from each sample was reverse transcribed. Transcript expression was assessed by qPCR on murine cDNAs diluted 1:25 performed using low ROX Takyon (Eurogentec) on a Viia 7 Real time PCR detection system (Life technologies). The exon-overlapping amplicons were amplified using the calibrated primers sets presented in Suppl Table 3. qPCRs were performed in triplicate for each sample. Results are expressed as 2^-ΔΔCt^ normalized on housekeeping gene expression (MmHPRT or RnPGK1 ; validated for a stable expression during cardiac remodelling) according to the Livak method.

### Oxygen consumption and extracellular acidification rate assessment

Oxygen consumption rate (OCR) and extracellular acidification rate (ECAR) were measured by using the Seahorse XF96 analyzer (Agilent, Santa Clara, CA, USA). Briefly, adult ventricular myocytes were seeded in Seahorse 96-well plates coated with laminin (10 000-15 000 cells/well; 6wells/condition) in M199 plating medium. After 1hr medium was replaced by 175 µL uncolored unbuffered DMEM pH 7,4 supplemented with 11mM Glucose and 1ng/L Insulin. OCR and ECAR values were assessed before after the injection of FCCP 1µM and Rotenone/Antimycin A 0,5µM with each step with 3 cycles of 3 min mixing/4 min measuring. Data were normalized by protein content in each well and expressed in pmol/min/µg protein or mpH/min/µg protein for OCR and ECAR, respectively.

### Transcriptomic analysis

AVM from β3AR-Tg (n=8) and WT littermates (n=8) mice were isolated 9 weeks post-TAC by cardiac Langendorff perfusion. Thereafter, AVM were plated for an hour before being processed for RNA extraction using Purelink RNA micro kit (thermofisher). After quality controls 500ng were processed for library preparation (TruSeq Stranded Total RNA LT Sample Prep Kit) after ribodepletion (Ribo-Zero). Sequencing was performed on Illumina NovaSeq 6000 - Paired end >60 millions reads. All sequencing data were analysed using the Automated Reproducible MOdular workflow for preprocessing and differential analysis of RNA-seq data (ARMOR) pipeline ^93^. In this pipeline, reads underwent a quality check using FastQC (Babraham Bioinformatics). Quantification and quality control results were summarised in a MultiQC report before being mapped using Salmon ^94^ to the transcriptome index which was built using all Ensembl cDNA sequences obtained in the Mus_musculus.GRCm39.cdna.all.fa file. Then, estimated transcript abundances from Salmon were imported into R using the tximeta package ^95^ and analysed for differential gene expression with edgeR ^96^. Over Representation Analysis was performed with the fGSEA Bioconductor package ^97^ on the mouse transcription factor recorded in the TRRUST v2 database ^23^. Raw and processed RNA-seq data were deposited and made publicly available on the Gene Expression Omnibus (GSE230859).

### Publicly available Single-nucleus RNA sequencing data re-analysis

Single-nucleus RNA sequencing data were analyzed by using a recently published dataset generated from patients with heart failure who recovered (versus patients who did not recover) left ventricular systolic function after left ventricular assist device implantation. ^22^. The corresponding dataset (GSE226314) available on the NCBI Gene Expression Ominibus (GEO) database was downloaded and analyzed with the Seurat 4.3.0 R package ^98^ to compute gene expression modulation in Cardiomyocyte cells as well as the WebGestaltR 0.4.5 ^99^ Bioconductor package to perform KEGG pathway over-representation analysis (ORA).

### Detection of reactive oxygen species

To assess the generation of intracellular ROS in cardiac myocytes, the peroxide-sensitive fluorescent probe 21,71-dichlorodihydrofluorescein (DCF) diacetate (Ref:C6827 Life Technologies) was used. AVM were seeded at density of 20 000 cells/well in 24-well plates coated with laminin. After 1hr plating, cells were incubated with 5µM H2-DCFDA for 30 min at 37°C. After incubation, AVM were washed 3 times with uncolored medium (113 mM NaCl, 4,7 mM KCl, 0,6 mM KH _2_PO_4_, 0,6 mM Na_2_HPO_4_, 1,2 mM MgSO_4_, 12 mM NaHCO3, 10 mM KHCO _3_, 30 mM Taurine, 10 mM BDM, 5,5 mM Glucose, 2nM Insulin, 1mM CaCl_2_, 0,1% BSA and 10 mM HEPES/NaOH, pH 7,46) and treated with ROS scavenger N-acetyl-cysteine 3mM. DCF fluorescence was detected at excitation and emission wavelengths of 494 and 523 nm, respectively, and measured in a fluorescence plate reader (Spectramax I3). The analysis of the data was made by using the Software SoftMax Pro. Data were normalized by protein content in each well and expressed as the increase in DCFDA/µg protein over a 30 min period in absence of treatment or the increase in DCFDA/µg protein over a 30 min period after treatment with 20µM H _2_O_2_ subtracted by the same condition in well pre-treated with ROS scavenger N-acetyl-cysteine to assess intracellular detoxification capacity of the myocytes.

### NRF2 activity determination

Nucleus were extracted from murine left ventricular tissues using the nuclear extraction kit (ab113474; abcam). Thereafter, obtained nuclear fractions were lyzed and protein were quantified. NRF2/NFE2L2 activity was quantified on similar quantity of nuclear extracts using the ELISA-based assay TransAM Nrf2 (50296; activemotif) measuring DNA binding activity of Nrf2 to adsorbed Antioxidant Response Elements (ARE) combined with an antibody-based detection by spectrophotometer.

### NRF2 immunostaining on isolated CM

NRVM were isolated as described above and cultured on chamber slides. After fixation in 4% formaldehyde, cells were permeabilized, blocked for aspecific staining and incubated with anti-NRF2 antibody (ab62352; 1:100) followed by AlexaFluor555-conjugated anti-rabbit antibody (A31572; thermofisher). Nuclei were counterstained with DAPI, slides were mounted with Dako fluorescence mounting medium and imaged using a Zeiss LSM800 confocal microscope. NRF2 nuclear and cytoplasmic stained area were quantified with Fiji using a thresholding method. To evaluate nuclear translocation, results were expressed as a repartition of the stained area between nuclear or cytosolic compartments. Total cell area was determined after manual delineation based on GFP expression. For each cellular isolation, 3-4 microscopic fields were acquired per condition and 6-10 cells analyzed per field.

### Statistical analysis

Results are expressed as mean ± SEM calculated from the average measurements obtained from each group of cells and mice. When normal distribution (tested by Kolmogorov-Smirnov test) was confirmed, raw data were analyzed using unpaired t-test or 2-way ANOVA followed by Sidak post-hoc test for multi-group comparisons. In absence of normal distribution, data were compared using nonparametric tests (Kruskall-Wallis followed by Dunn correction for multiple comparisons or Mann–Whitney). Statistical significance was accepted at the level of P<0,05.

## Acknowledgements

We thank D. Thibou and E. Vandenhooft for handling the β3AR-Tg mouse line.

## Declaration of interests

The authors declare no competing interests.

**Suppl Fig S1 -Cardiac β3AR protects against myocardial hypertrophy without compromising myocardial function –** Percentage of **(A)** ejection fraction and **(B)** fractional shortening in β3AR-Tg and WT littermates before and after TAC for 9 weeks. Results are expressed as means ± SEM with *P < 0,05 calculated by Two-way Anova and corrected for multiple comparisons with Sidak’s test.

**Suppl Fig S2 – Unchanged circulating metabolites – (A-C)** Volcano plot displaying unbiased serum metabolomics **(A)** from 9-week TAC WT compared with control, non-operated WT mice by reverse phase LC/MS (M1) or HILIC LC/MS (M2) in negative (neg) and positive (pos) ionization mode; **(B)** from 9-week TAC β3AR-Tg and 9-week TAC WT mice by HILIC LC/MS **(C)** from Control β3AR-Tg and Control WT mice by reverse phase LC/MS (M1) or HILIC LC/MS (M2). Numbers of detected and plotted signals are reported in Methods. P value corrected by False discovery rate approach < 0,05

**Suppl Fig S3 – Unchanged myocardial lipid uptake levels – (A)** 18-^18^F-Fluoro-4-Thia-Oleate assessment of myocardial fatty acid uptake *in vivo* by PET (expressed as standardized uptake value) and **(B)** Increase in ^14^C-Palmitate without or **(C)** following insulin-treatment in AVM isolated from 9-week TAC or Control β3AR-Tg or wild-type mice. **(D)** Fatty acids transporter CD36 in AVM from 9-week TAC or Control β3AR-Tg or wild-type mice. Results are expressed as means ± SEM with *P < 0,05 calculated by Two-way Anova and corrected for multiple comparisons with Sidak’s test.

**Suppl Fig S4 ‒ Cardiac β3AR does not alter lactate production -** Extracellular acidification rate measured by Seahorse analyzer in AVM isolated from 9-week TAC or Control β3AR-Tg or wild-type mice. Results are expressed as means ± SEM with *P < 0,05 calculated by Two-way Anova corrected for multiple comparisons with Sidak’s test.

**Suppl Fig S5 – Recovery of cardiac function in patients is paralleled with expression of transcripts corresponding to pathways associated with NRF2 activation** - Enrichment ratio of KEGG pathways computed with an over-representation analysis (ORA) of differentially expressed genes in nuclear extracts from cardiac myocytes isolated from biopsies obtained in patients who favorably responded or not to the implantation of a Left Ventricular Assist Device (LVAD). The results were computed from a publicly available database reporting transcripts abundance determined by single nucleus RNA-sequencing and published in ^22^ Differentially enriched pathways for transcripts of the PPP, insulin resistance and cGMP-PKG pathways (activated by nitric oxide) are highlighted in bold characters.

