## Supplemental figures for "A NRF2/β3-adrenoreceptor axis drives a sustained antioxidant and metabolic rewiring through the pentose-phosphate pathway to alleviate cardiac stress"

#### Supplemental Figure S1

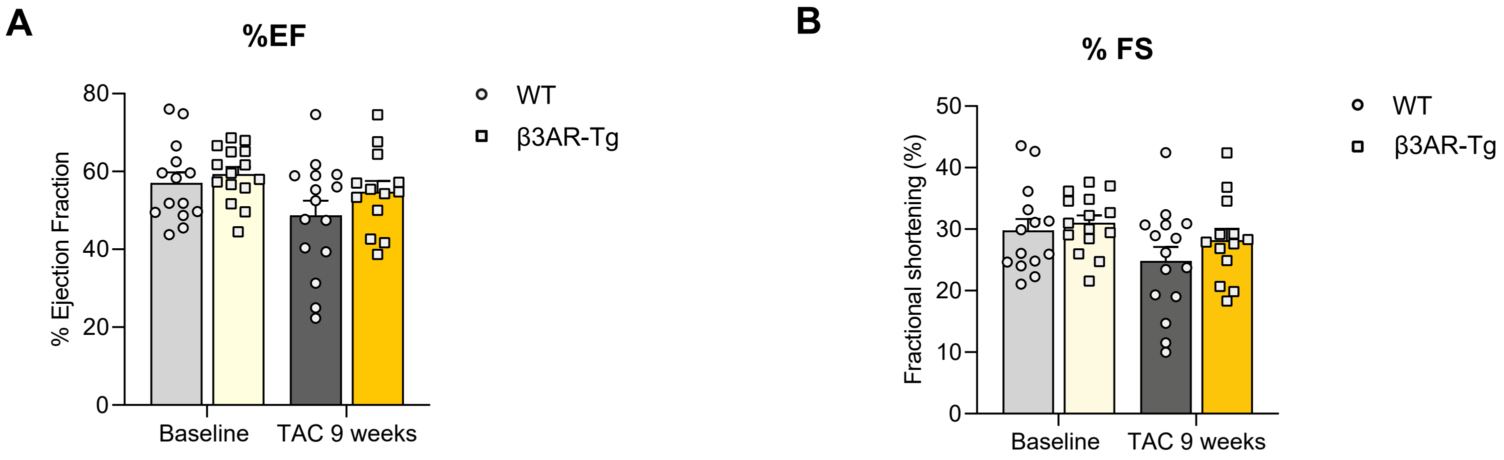

#### Supplemental Figure S2

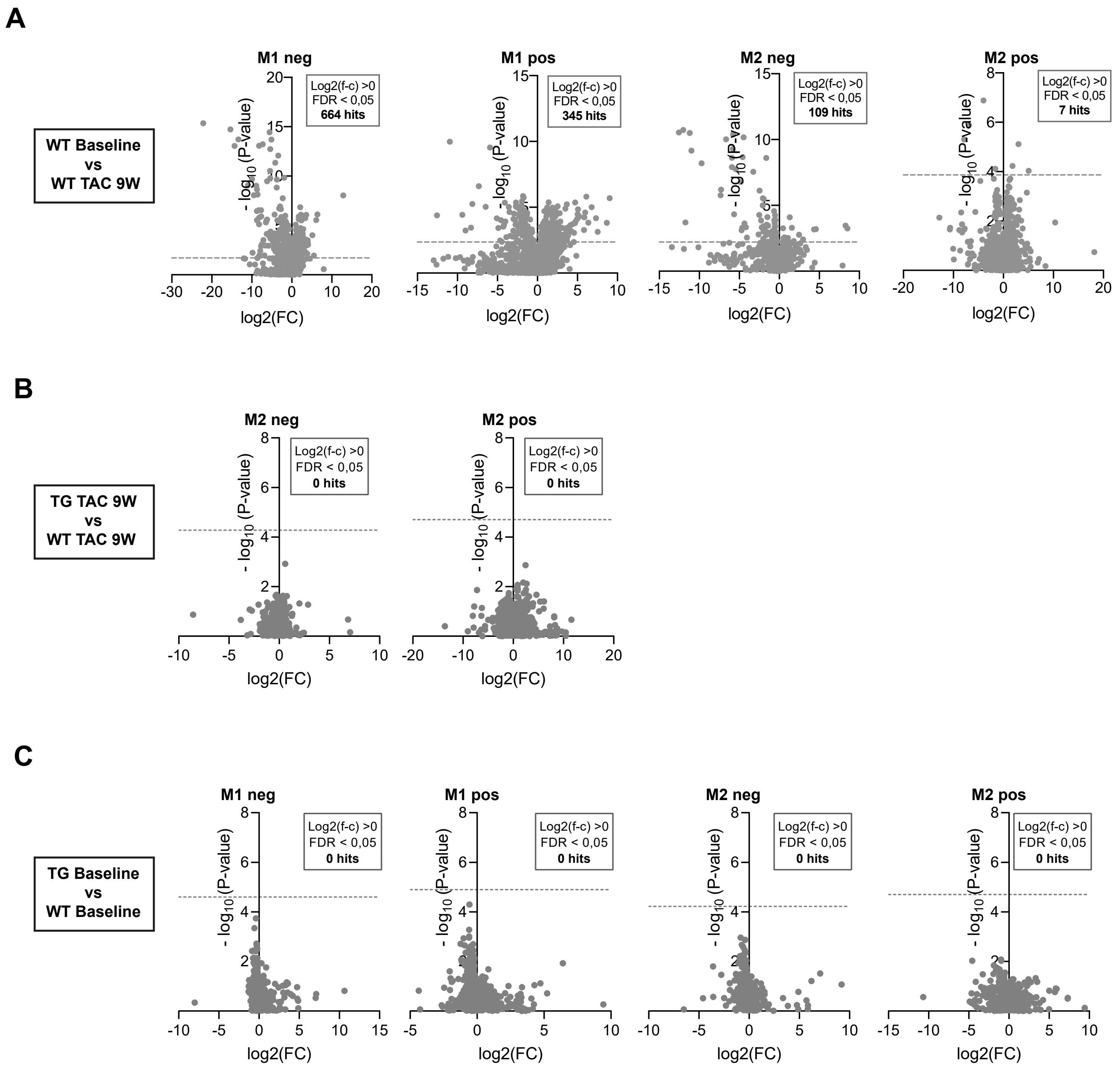

### Supplemental Figure S3

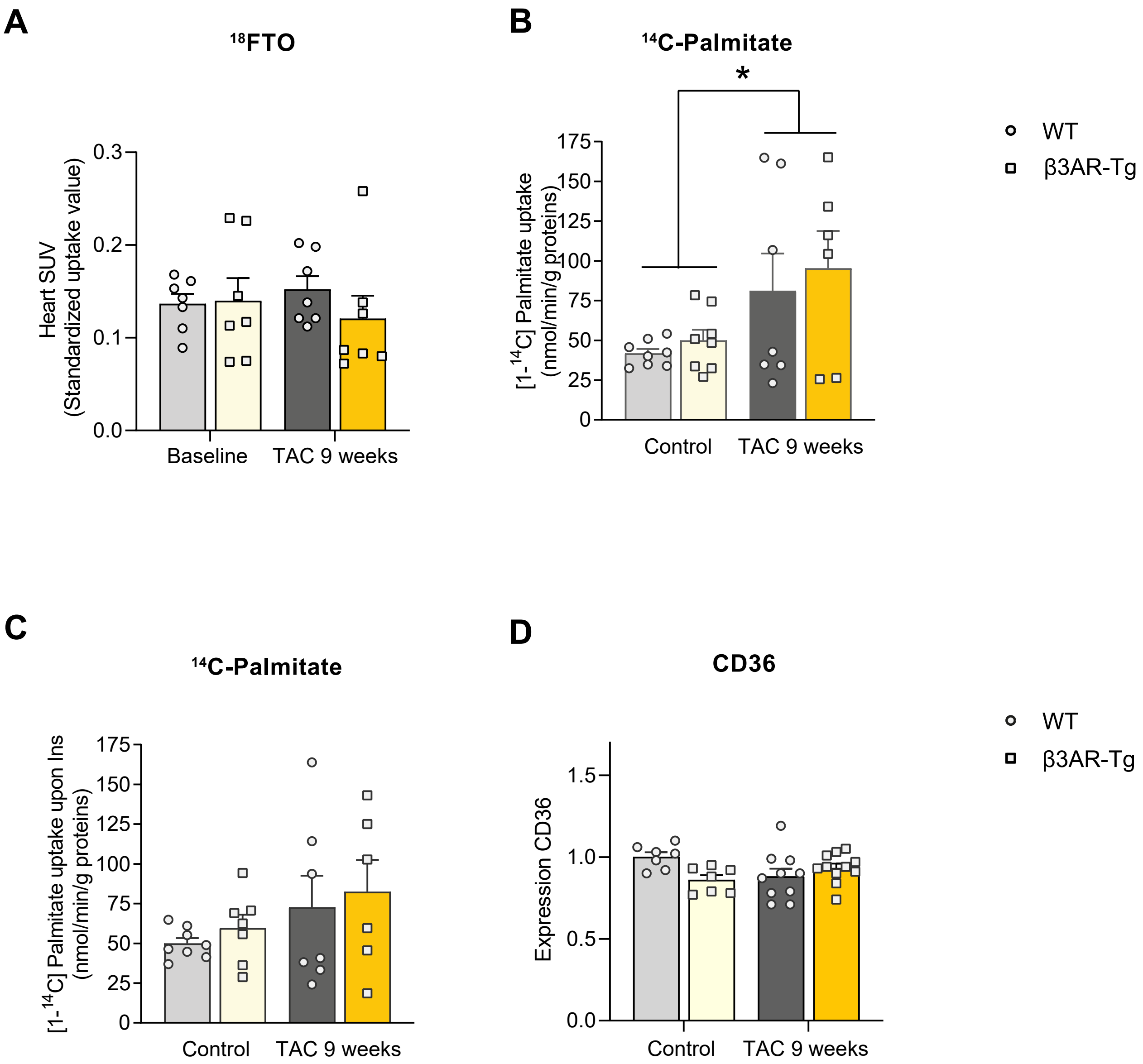

**Supplemental Figure S4**

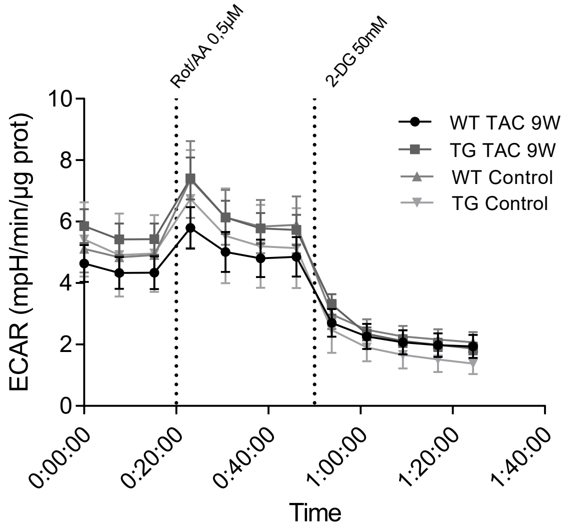

### Supplemental Figure S5

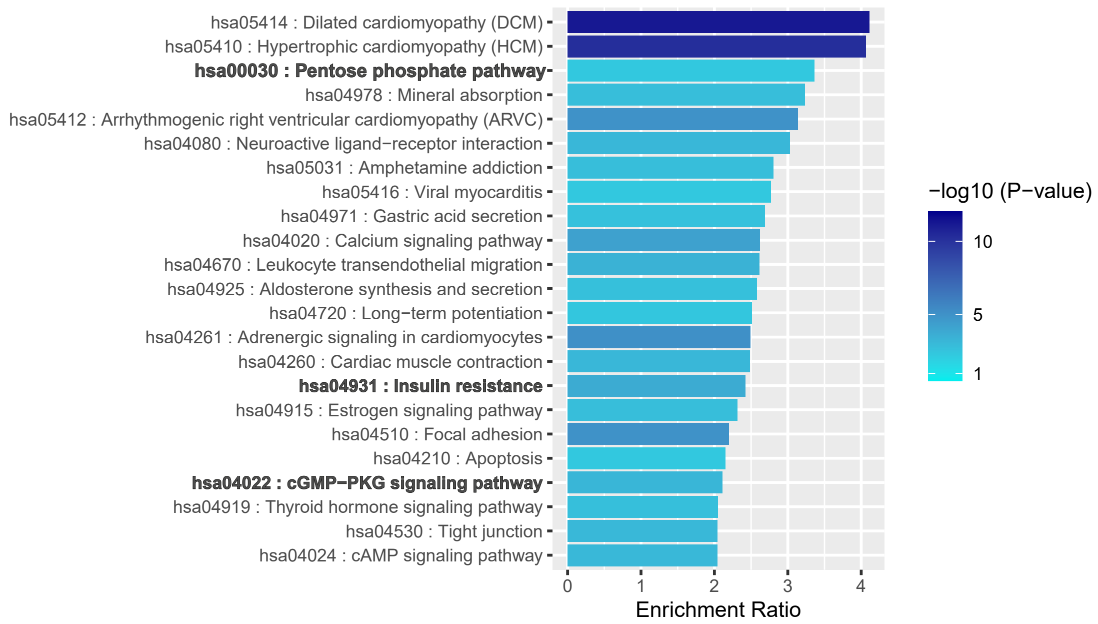
