## Supplemental Tables for "A NRF2/β3-adrenoreceptor axis drives a sustained antioxidant and metabolic rewiring through the pentose-phosphate pathway to alleviate cardiac stress"

**Supplemental Table 1 : Echocadiographic measurements**

| | Wild type Baseline | B3-Tg $^{+/-}$ Baseline | Wild type TAC 9W | B3-Tg $^{+/-}$ TAC 9W |
|---|---|---|---|---|
| **Body weight (g)** | 24,9 ± 0,7 | 25,7 ± 0,5 | 26,3 ± 0,7 | 28,4 ± 0,8 |
| **Aortic velocity (ms)** | - | - | 3,4 ± 0,4 | 3,4 ± 0,3 |
| **Heart Rate (bpm)** | 436 ± 10 | 483 ± 11 | 462 ± 13 | 511 ± 11 |
| **LVIDd (mm)** | 3,72 ± 0,07 | 3,67 ± 0,05 | 4,00 ± 0,11 | 3,67 ± 0,06 |
| **LVIDs (mm)** | 2,63 ± 0,10 | 2,53 ± 0,07 | 3,03 ± 0,16 | 2,64 ± 0,09 |
| **LVPWd (mm)** | 0,69 ± 0,02 | 0,69 ± 0,02 | 0,99 ± 0,04 | 0,94 ± 0,03 |
| **LVPWs (mm)** | 1,05 ± 0,02 | 1,06 ± 0,02 | 1,32 ± 0,03 | 1,30 ± 0,02 |
| **IVSd (mm)** | 0,64 ± 0,01 | 0,64 ± 0,01 | 1,00 ± 0,04 | 0,91 ± 0,04 |
| **IVSs (mm)** | 1,00 ± 0,02 | 1,03 ± 0,01 | 1,31 ± 0,03 | 1,24 ± 0,02 |
| **EF (%)** | 57,0 ± 2,7 | 59,3 ± 1,8 | 48,7 ± 3,8 | 54,8 ± 2,8 |
| **FS (%)** | 29,8 ± 1,9 | 31,0 ± 1,2 | 24,8 ± 2,3 | 28,2 ± 1,9 |
| **Stroke Vol (µL)** | 33,2 ± 1,6 | 33,4 ± 1,3 | 32,2 ± 2,1 | 31,1 ± 1,4 |
| **LV vol s (µL)** | 59,4 ± 2,7 | 57,2 ± 1,8 | 69,8 ± 4,7 | 57,7 ± 2,0 |
| **LV vol d (µL)** | 26,1 ± 2,7 | 23,7 ± 1,6 | 37,6 ± 4,8 | 26,6 ± 2,0 |

LVID, left ventricle internal diameter in diastole (LVIDd) or systole (LVIDs); LVPW, posterior wall thickness in diastole (LVPWd) or systole (LVPWs); IVS, interventricular septum thickness in diastole (IVSd) or systole (IVSs); EF, ejection fraction; FS, fractional shortening; LV vol, left ventricular volume in diastole (LV vol d) or systole (LV vol d).

**Supplemental Table 2 : Characteristics of targeted metabolites detection**

| Metabolite | Retention time | m/z | Mass | Formula | ID | Category |
|---|---|---|---|---|---|---|
| Methionine_sulfoxide | 1.51 | 164.03869 | 165.0459665 | C5H11NO3S | HMDB02005 | Ion Pairing |
| D-Sedoheptulose_7-phosphate | 9.88 | 289.03301 | 290.0402865 | C7H15O10P | HMDB01068 | Ion Pairing |
| Fructose_6-phosphate | 10.15 | 259.02244 | 260.0297165 | C6H13O9P | HMDB00124 | Ion Pairing |
| Pentose_phosphate | 10.23 | 229.01188 | 230.0191565 | C5H11O8P | HMDB01548 | Ion Pairing |
| Dihydroxyacetone_phosphate | 10.7 | 168.99075 | 169.9980265 | C3H7O6P | HMDB01473 | Ion Pairing |
| Glutathione | 11.03 | 306.07653 | 307.0838065 | C10H17N3O6S | HMDB00125 | Ion Pairing |
| CMP | 11.34 | 322.04457 | 323.0518465 | C9H14N3O8P | HMDB00095 | Ion Pairing |
| UMP | 11.96 | 323.02859 | 324.0358665 | C9H13N2O9P | HMDB00288 | Ion Pairing |
| GMP | 12.14 | 362.05072 | 363.0579965 | C10H14N5O8P | HMDB01397 | Ion Pairing |
| AMP | 13.57 | 346.05581 | 347.0630865 | C10H14N5O7P | HMDB00045 | Ion Pairing |
| Oxidized_glutathione | 13.88 | 611.14469 | 612.1519665 | C20H32N6O12S2 | HMDB03337 | Ion Pairing |
| Uridine_diphosphate_glucose | 14.51 | 565.04774 | 566.0550165 | C15H24N2O17P2 | HMDB00286 | Ion Pairing |
| Phosphoenolpyruvic_acid | 16.42 | 166.9751 | 167.9823765 | C3H5O6P | HMDB00263 | Ion Pairing |
| Fructose 1,6-bisphosphate | 16.73 | 338.98877 | 339.9960465 | C6H14O12P2 | HMDB01058 | Ion Pairing |
| 3-Phosphoglyceric_acid | 17.08 | 184.98566 | 185.9929365 | C3H7O7P | HMDB00807 | Ion Pairing |
| Uridine_diphosphate-N-acetylglucosamine | 14.81 | 606.07429 | 607.0815665 | C17H27N3O17P2 | HMDB00290 | Ion Pairing |
| NADPH | 12.85 | 744.08383 | 745.0911065 | C21H30N7O17P3 | HMDB00221 | HILIC |
| NADP | 14.29 | 742.06818 | 743.0754565 | C21H28N7O17P3 | HMDB00217 | HILIC |

**Supplemental Table 3 : Primers sequences**

| Target | Forward | Reverse |
|---|---|---|
| Mm_HPRT | TTG-GAT-ACA-GGC-CAG-ACT-TTG | CGT-GAT-TCA-AAT-CCC-TGA-AG |
| Mm_PGD | CCC-CAC-ATC-AAG-GCG-ATC-TT | TCA-TCT-CCC-ACC-CAG-TCA-CA |
| Mm_SLC2A1 | ATC-ATC-GGT-TAC-TGC-GG | ACGC-CAA-ACA-CCT-GGG-CAA-TA |
| Mm_SLC2A4 | GGG-TGG-CAT-GAT-CTC-TTC-CTT-T | GAG-TAG-GCG_CCA-ATG-AGG-AAC |
| Mm_NPPA | TCGTCTTGGCCTTTTGGCT | TCCAGGTGGTCTAGCAGGTTCT |
| Mm_NPPB | GCCAGTCTCCAGAGCAATTC | AGCTGTCTCTGGGCCATTT |
| Mm_MYH6 | TGTGGTGCCTCGTTCCA | TTTCGGAGGTACTGGGCTG |
| Mm_MFN1 | ACT-CAG-TAA-ACG-TGG-CAG-CA | TCC-TCC-GTG-ACC-TCC-TTG-AT |
| Mm_CD36 | TGG-CCA-AGC-TAT-TGC-GAC-AT | TTC-AGA-TCC-GAA-CAC-AGC-GT |
| Mm_OPA1 | TAC-TGT-TAG-CCC-CGA-GAC-CA | GAT-GAC-ACC-AGG-CAA-GTC-CA |
| Mm_PHKG | CAT-CCT-GCA-GAA-GGT-CTC-GG | CCC-TCT-CTT-CAT-CAG-ATC-AAA-TAC-C |
| Mm_PHKA | TCC-AAC-ACT-GCC-AGT-CTA-TCC | TCC-ACC-AAA-TGG-CTG-TCC-TC |
| Mm_Ppargc1a | AAA-CTT-GCT-AGC-GGT-TCT-CA | TGG-CTG-GTG-CCA-GTA-AGA-G |
| Mm_TFAM | TCC-ACA-GAA-CAG-CTA-CCC-AA | CCA-CAG-GGC-TGC-AAT-TTT-CC |
| Mm_NFE2L2 | GGA-CAT-GGA-GCA-AGT-TTG-GC | CAG-CGG-TAG-TAT-CAG-CCA-GC |
| Mm_MFN2 | ACT-CAG-TAA-ACG-TGG-CAG-CA | TCC-TCC-GTG-ACC-TCC-TTG-AT |
| Mm_Taldo | TTA-TCA-TCA-ACC-TGG-GAG-GG | GCG-AAG-GAG-AAA-AGC-AGT-GT |
| Mm_Myh7 | ACC-AAC-CTG-TCC-AAG-TTC-CG | ACT-CCT-CAT-TCA-GGC-CCT-TG |
| Mm_GPX1 | ATG-TGT-GCT-GCT-CGG-CTC | AGA-GAG-ACG-CGA-CAT-TCT-CAA |
| Mm_G6PD | GGT-CGT-GGG-GGC-TAT-TTT-GA | GAA-CCT-GTA-GTG-GCA-GGC-TT |
| Rn_PGK1 | GAA-GGG-AAG-GGA-AAA-GAT-GC | AAA-TCC-ACC-AGC-CTT-CTG-TG |
| Rn_NFE2L2 | CAC-ATC-CAG-ACA-GAC-ACC-AGT | CTA-CAA-ATG-GGA-ATG-TCT-CTG-C |
| Rn_NPPA | AGG-ATT-GGA-GCC-CAG-AGC-GGA-CTA-GG | TGA-TAG-ATG-AAG-ACA-GGA-AGC-TGC |
| Rn_NPPB | GCC-AGT-CTC-CAG-AAC-AAT-CC | AGC-TGT-CTC-TGA-GCC-ATT-T |
